# Correlates of protection against African swine fever virus identified by a systems immunology approach

**DOI:** 10.1101/2025.05.25.655978

**Authors:** Kirill Lotonin, Francisco Brito, Kemal Mehinagic, Obdulio García-Nicolás, Matthias Liniger, Noelle Donzé, Sylvie Python, Stephanie Talker, Tosca Ploegaert, Nicolas Ruggli, Charaf Benarafa, Artur Summerfield

## Abstract

African swine fever virus (ASFV) causes a fatal hemorrhagic disease in domestic pigs and wild boars, which poses severe threats to the global pork industry. Despite the promise of live attenuated vaccines (LAVs), their narrow margin between efficacy and residual virulence presents major safety challenges. This study bridges a critical knowledge gap in ASF vaccinology by identifying innate and adaptive correlates of protection. This was achieved by using an established model with two groups of pigs differing in baseline immunological status (farm and specific pathogen-free [SPF]). The animals were immunized with an attenuated ASFV strain and subsequently challenged with a related, highly virulent genotype II strain. By applying a systems immunology approach, we correlated kinetic data, including serum cytokines, blood transcription modules (BTMs), T-cell responses, and antibody levels, with clinical outcomes to track protective and detrimental immune responses to the virus over time. Key innate correlates of protection included early and sustained IFN-α response, activation of antigen presentation BTMs, and controlled IL-8 levels during immunization. Lower baseline immune activation observed in SPF pigs in steady state was linked to increased protection. Adaptive correlates encompassed cell cycle, plasma cell, and T-cell BTM responses lasting until day 15 post-immunization. Consequently, an effective response from ASFV-specific T_h_ cells prior to challenge indicated protection. After the challenge, an early IFN-α response, along with low levels of pro-inflammatory cytokines and a strong induction of memory T_h_ and T_c_ cells, correlated with improved clinical outcomes. The model highlights the critical role of host-specific factors in vaccine efficacy and provides a valuable framework for optimizing ASFV vaccine design while distinguishing between protective and detrimental immune responses.

## Introduction

African swine fever virus (ASFV), a large and complex DNA virus in the *Asfarviridae* family, is the causative agent of a severe hemorrhagic disease affecting both domestic and wild pigs of the *Sus* genus, with high mortality rates. The disease caused by this macrophage-tropic virus is characterized by dramatic immune dysregulation and cytokine storm (Gaudreault et al., 2020; Li et al., 2022). Since its introduction to Georgia in 2007, ASFV genotype II has expanded extensively across Eurasia, significantly impacting global pig populations (Wang et al., 2018; Sauter-Louis et al., 2021; Mahanta et al., 2024). The virus is highly resistant in the environment and continues to spread in suid populations, especially in areas with poor biosecurity measures.

At present, two live attenuated vaccines (LAVs) are registered and used only in Vietnam (Fan et al., 2024). Only LAVs have been shown to induce robust protection against homologous challenge, but their use remains associated with safety concerns, which has hindered their widespread application (Gladue et al., 2022; Urbano and Ferreira, 2022). A multitude of LAV candidates has been developed by deleting non-essential genes, followed by testing for virulence and protective capacity in vaccination-challenge experiments. Since then, a trade-off between safety and the induction of protective immunity has been a major problem. For this reason, it is crucial to identify candidates that are effective and safe across a broad range of conditions (Chu et al., 2024).

The lack of knowledge on immunological mechanisms underlying both protection and disease aggravation remains a major obstacle in addressing the issues mentioned above. The role of T cells and antibodies in protection against genotype I strains was experimentally demonstrated by CD8α^+^ T cell depletion (Oura et al., 2005) and the adoptive transfer of immune sera (Onisk et al., 1994). Although several studies have emphasized the importance of T-cell responses following infection or vaccination (Takamatsu et al., 2013; Schäfer et al., 2022; Pedrera et al., 2024), the mathematical correlation of these responses with protection after LAV application has not been established, and we still lack reliable readouts that indicate protective immunity (Wang et al., 2023).

Considering the immunopathogenesis of ASF, it is also critical to identify protective and detrimental innate immune responses. They often appear to lack regulation during ASFV infection that can result in a deleterious cytokine storm (Ayanwale et al., 2022; Franzoni et al., 2023). Moreover, it is essential to explore the interaction between innate and adaptive immunity to better understand the proper induction of the latter (Bosch-Camós et al., 2022).

Based on these knowledge gaps, the present study was initiated to identify innate and adaptive correlates of protection. We combined clinical, cytokine, adaptive immune profiling and transcriptomic kinetic datasets to enable a systems immunology approach to dissect protective and detrimental immune responses following immunization and challenge with two ASFV strains of genotype II with differing virulence (Lacasta et al., 2015; Sereda et al., 2022; Lin et al., 2022; Gonçalves et al., 2024). To obtain the required variation in disease development and immune responses, we immunized farm and SPF pigs with the attenuated Estonia 2014 strain. SPF pigs have previously been reported to be more resilient to Estonia 2014 infection due to a lower inflammatory response and were also better protected against the virulent Armenia 2008 challenge upon immunization (Radulovic et al., 2022, 2025). To capture immunologically relevant gene interaction networks, we employed blood transcription modules, or BTMs (Li et al., 2014), that were successfully implemented for studying immune responses to vaccines in humans (Kazmin et al., 2017; Hagan et al., 2022) and animals (Braun et al., 2018; Matthijs et al., 2019; Bocard et al., 2021).

Building on the importance of baseline immunity for vaccination (Fourati et al., 2022; Toptygina et al., 2023), we mimicked variations in baseline immune activation using breed- and age-matched farm and SPF pigs. Our data provide a temporally resolved model of protective and detrimental innate and adaptive immune responses, based on multiple immunological parameters, and offer a framework for assessing ASFV vaccine candidates.

## Results

### Improved development of protective immunity in SPF compared to farm pigs

To define immune correlates of protection in immunized pigs against virulent challenge infection, we used the previously established model, in which the baseline immunological status impacts ASF progression, inflammatory responses, as well as the ability to develop protective immunity (Radulovic et al., 2022, 2025). Accordingly, two groups of pigs - farm and SPF - were first immunized with the attenuated Estonia 2014 strain and monitored in terms of clinical signs, viral loads, systemic cytokine responses, and transcriptome profiles. The pigs were also tested for virus-specific T-cell responses before and after challenge with the highly virulent Armenia 2008 strain (Fig. 1A).

**Fig. 1.**
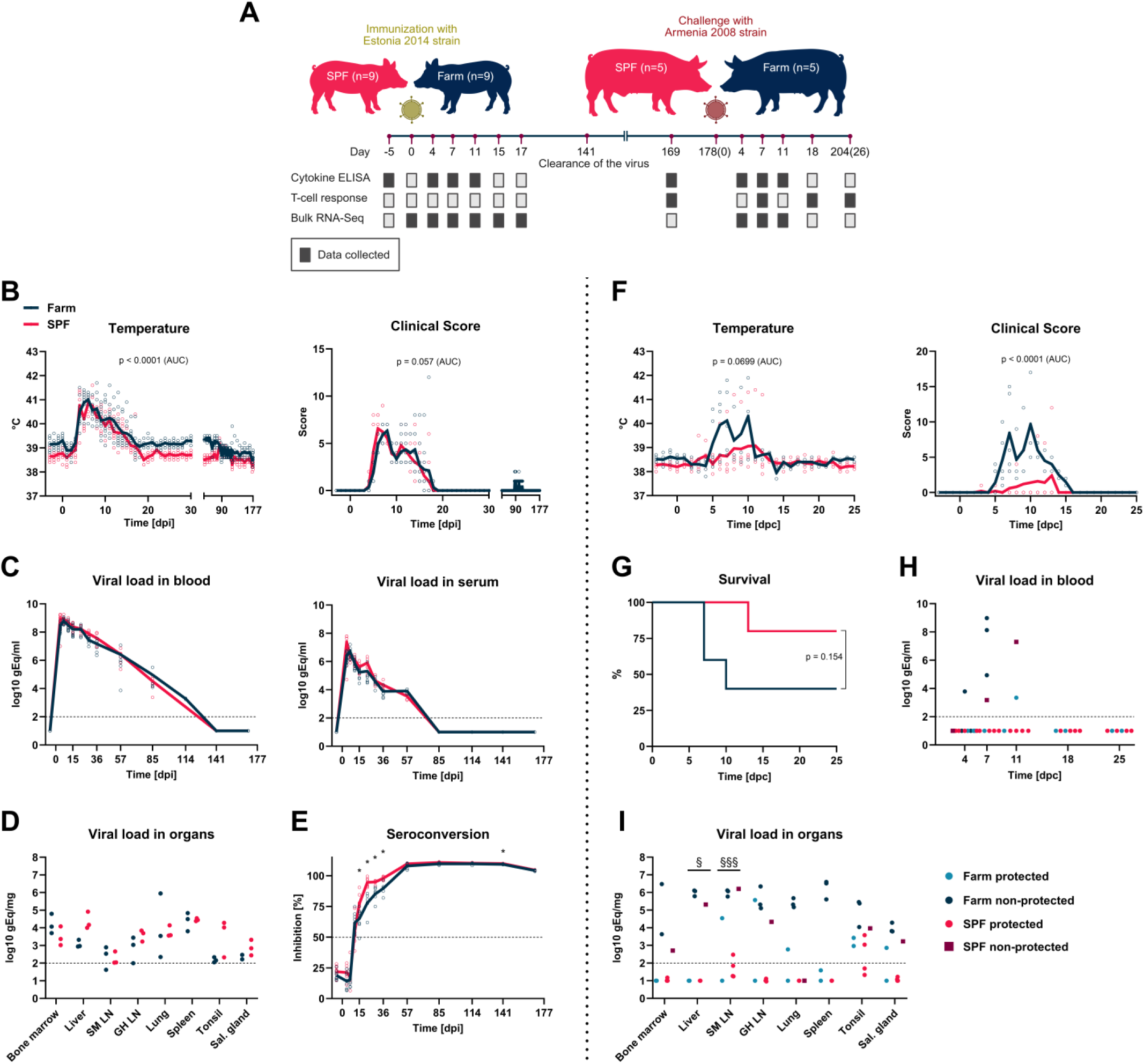
Outcomes of ASFV immunization and challenge in SPF and farm pigs. (A) Overview of experiment layout marking time points of sample collection and performed immunological readouts (dark rectangles indicate collected data). (B, F) Rectal temperatures and clinical scores were recorded daily after immunization with the attenuated Estonia 2014 strain and after challenge with the highly virulent Armenia 2008 strain. (C, H) Viral DNA levels in blood and serum were measured by qPCR. (D, I) Organs were collected at 17 dpi (D) and at 7, 10, 13 and 26 dpc (I) for quantification of viral DNA levels (SM LN – submandibular lymph node; GH LN – gastrohepatic lymph node). (E) Seroconversion was analyzed by competitive ELISA. For B, C, E, and F, the data points represent values from individual animals and the lines show means of the groups. For D, H, and I, the data points represent values for individual animals.There were n = 9 pigs per group in B, C, and E, and n = 3 in D. In F and G, there were n = 5 pigs per group at 0 dpc. In H, at 4 and 7 dpc, n = 5 pigs in farm and SPF groups; at 11 dpc, n = 2 in farm group and n = 5 in SPF group; and at 18 and 25 dpc, n = 2 in farm group and n = 4 in SPF group. In I, n = 5 pigs in farm and SPF groups. For B and F, the differences in temperature and clinical scores between two groups were analyzed by comparing areas under curves (AUC) with unpaired t-test. For C, D, and E, the differences between farm and SPF groups were analyzed at each time point by unpaired t-test with Holm-Šídák’s correction for multiple comparisons; *p < 0.05. For G, the survival rates in two groups were compared by Log-rank test. For H and I, the differences in viral loads between protected and non-protected animals were analyzed by unpaired t-test with Holm-Šídák’s correction for multiple comparisons; ^§^p < 0.05; ^§§§^p < 0.001.

After Estonia 2014 infection, both farm and SPF pigs developed similar clinical symptoms and recovered completely by 20 days post-infection (dpi; Fig. 1B). Also, the viremia was comparable in both groups, reaching a peak in the first week and being cleared at 141 dpi (Fig. 1C). Three pigs from each group were sacrificed at 17 dpi, and viral loads were examined in collected organs, again with no statistical difference identified between farm and SPF pigs (Fig. 1D). Furthermore, both groups seroconverted with a similar kinetic, although the competition ELISA indicated higher anti-p72 antibody titers between 15 and 36 dpi for SPF pigs (Fig. 1E). Both farm and SPF pigs showed a decrease in the number of white blood cells and platelets early after the infection, and by 22 dpi the indices returned to the baseline level. As previously reported by Radulovic et al. (2022, 2025), the counts of monocytes and neutrophils were generally higher in farm animals. In contrast to that, red blood cell levels were higher in SPF pigs, and showed a slow decline in both groups after immunization (Supplementary Fig. 1).

Following challenge with the Armenia 2008 strain at 178 dpi (0 days post-challenge; dpc), farm pigs demonstrated a more severe disease course. Three pigs reached the discontinuation criteria specified in the animal experimentation license and had to be sacrificed at 7 and 10 dpc. The other two farm pigs developed clinical signs but recovered by 16 dpc. In contrast, four of the five SPF pigs showed no clinical symptoms, while one animal developed severe ASF and had to be sacrificed at 13 dpc (Fig. 1F, G). Non-protected pigs had detectable viral loads in blood and serum, peaking at 7 dpc (Fig. 1H and Supplementary Fig. 2). High viral loads were also mainly observed in organs from non-protected animals, most prominently in liver and lymph nodes (Fig. 1I). Moreover, only in non-protected animals the challenge infection caused a decrease in blood lymphocytes, neutrophils and platelets at 7-11 dpc (Supplementary Fig. 3).

### Cytokine correlates of protection: role of IFN-α and controlled inflammatory response

Given that viral loads during immunization with the Estonia 2014 strain were similar between the groups and did not explain the different outcomes of the challenge infection, we hypothesized a role of the immune response in defining protection. As cytokines are primary regulators of innate and adaptive immune responses, and are induced systemically following ASFV infection, we investigated their differential dynamics in two groups of pigs, as well as their correlation with protection. Both farm and SPF pigs had a peak of IFN-α response at 4 dpi that persisted above baseline at 7 and 11 dpi. When compared to the SPF group, the farm pigs had elevated baseline levels of IL-1β and IL-8, as well as stronger responses for IL-8 at 4 and 7 dpi, and for IFN-γ and TNF at 7 dpi (Fig. 2A). Next, we calculated the correlation of cytokine responses with protection defined as area under the curve (AUC) for clinicals scores (Supplementary Fig. 4). This identified IFN-α at 7 dpi as a potential factor of improved clinical outcome, while higher levels of IL-8 at 7 dpi correlated positively with higher clinical scores (Fig. 2B and Supplementary Fig. 5).

**Fig. 2.**
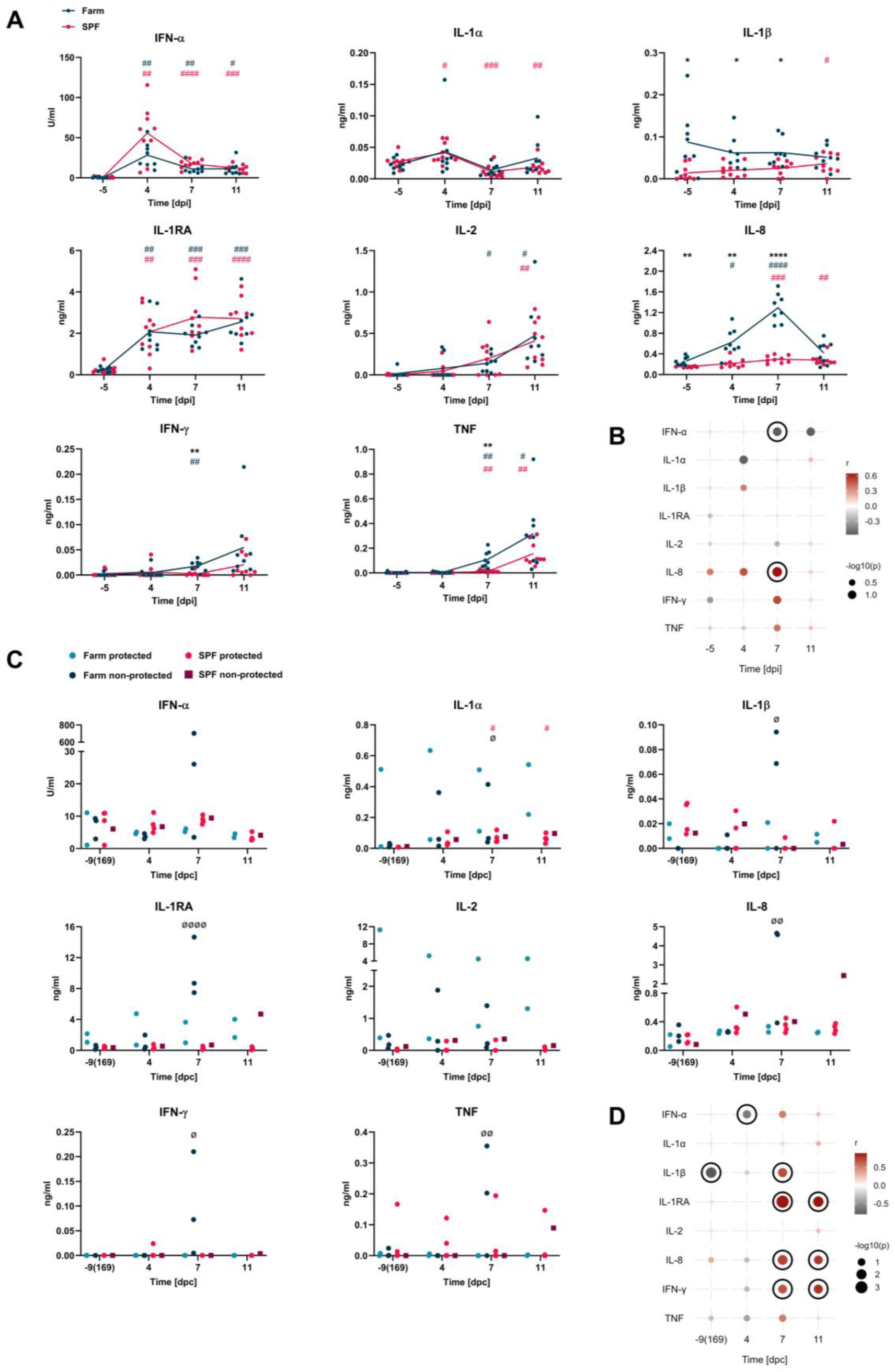
Cytokine responses following immunization and challenge, and their correlation with protection. (A, C) Serum cytokine levels were measured by ELISA. (B, D) Dot plots show correlations between cytokine levels and clinical scores after challenge (r – correlation coefficient; dark red indicates a negative correlation with protection, which reflects poor clinical outcomes, while gray indicates a positive correlation with protection; correlations with p < 0.1 are outlined with black circles). For A, the data points represent values for individual animals and the lines show means of the groups. For C, the data points represent values for individual animals. In A, there were n = 8 farm pigs and n = 9 SPF pigs, and in C, at -9, 4 and 7 dpc, n = 5 pigs in farm and SPF groups, and at 11 dpc, n = 2 in farm group and n = 5 in SPF group. In A, the differences to baseline measurements (day -5) for farm and SPF pigs were analyzed by two-way ANOVA with Dunnett’s multiple comparisons test (blue hashtags - farm group, pink hashtags - SPF group); ^#^p < 0.05; ^##^p < 0.01; ^###^p < 0.001; ^####^p < 0.0001. Differences between farm and SPF groups were analyzed at each time point by unpaired t-test with Holm-Šidák’s correction for multiple comparisons; *p < 0.05; **p < 0.01; ****p < 0.0001. In C, the differences to baseline measurements (day -9) for farm, SPF, protected and non-protected groups were analyzed by mixed-effects analysis with Dunnett’s multiple comparisons test (pink hashtags – SPF group, Ø – non-protected animals); ^#^p < 0.05; ^Ø^p < 0.05; ^ØØ^p < 0.01; ^ØØØØ^p < 0.0001. Differences between farm and SPF groups, as well as protected and non-protected animals, were analyzed at each time point by unpaired t-test with Holm-Šidák’s correction for multiple comparisons.

After challenge with the highly virulent Armenia 2008 strain, cytokine levels were highly variable, but indicated significantly higher levels of IL-1β, IL-1RA, IL-8, IFN-γ and TNF at 7 dpc in non-protected farm animals. Interestingly, IL-1β was not detectable before challenge in the farm animals with the most severe clinical outcomes (Fig. 2C). Correlation of the cytokine data with clinical scores demonstrated that sustained levels of IL-1β before the challenge, as well as early IFN-α upregulation at 4 dpc were beneficial for the animals. On the contrary, the levels of IL-1β, IL-1RA, IL-8, and IFN-γ at 7, and occasionally at 11 dpc, correlated negatively with protection (Fig. 2D and Supplementary Fig. 6).

### T_h_-cell response represents the main correlate of protection

To identify T-cell responses correlating with protection against virulent ASFV infection, PBMCs collected before and after the challenge were restimulated with the virus *in vitro*. IFN-γ ELISpot data demonstrated presence of virus-specific cytokine-secreting lymphocytes before challenge in all animals, but with a high level of variation (Fig. 3A). Interestingly, IFN-γ secretion became undetectable or low in all non-protected pigs at 7 dpc. Next, we examined effector cytokine production by specific immune cell subsets using flow cytometry (Fig. 3B). The numbers of TNF^+^ and TNF^+^/IFN-γ^+^ virus-specific CD4^+^CD8α^+^ double-positive (DP) T_h_ cells were higher in SPF pigs before the challenge. Furthermore, at 7 dpc, only protected animals showed TNF/IFN-γ and IFN-γ recall responses from this cell population. Protected and non-protected pigs could also be distinguished by comparing TNF/IFN-γ and IFN-γ cytokine responses from CD4^+^CD8α^-^ single-positive (SP) T_h_ cells and CD8α^+^CD8ꞵ^+^ T_c_ cells at 7 dpc. Besides that, protected farm and SPF pigs had higher levels of IFN-γ^+^ effector γδ T cells at 7 dpc. Notably, survived farm pigs had a substantial production of IFN-γ by NK cells at 7 dpc (Supplementary Fig. 8). Correlation of effector cytokine responses with clinical scores before and after challenge demonstrated that activation of DP and SP T_h_ cells, as well as T_c_ cells, correlated with protection. Nevertheless, only responses from SP and DP T_h_ cells correlated with improved clinical outcomes when measured before challenge (Fig. 3C and Supplementary Fig. 9, 10).

**Fig. 3.**
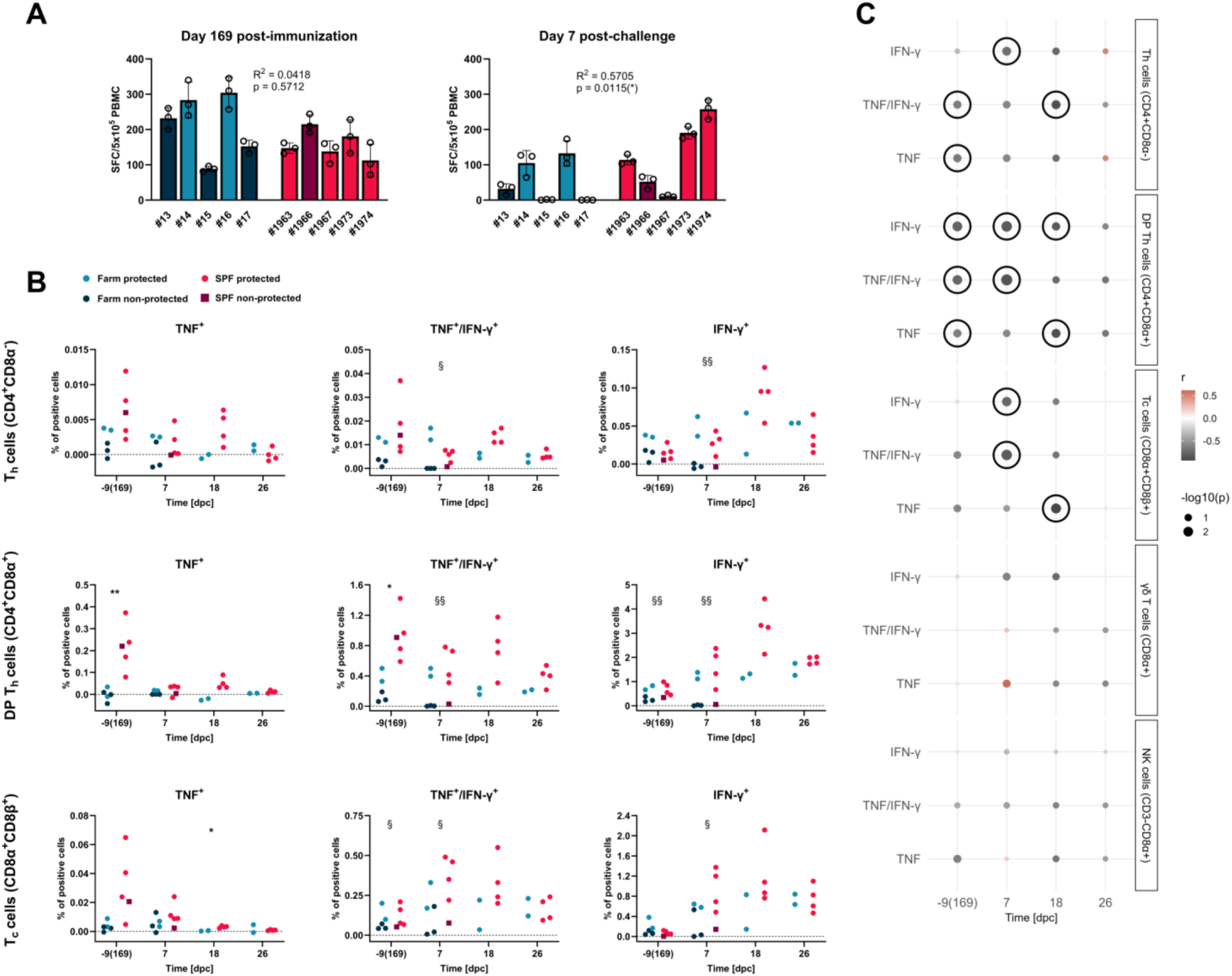
Cellular immune responses before and after challenge and their correlation with protection. (A) IFN-γ release from freshly collected PBMCs upon ASFV restimulation was measured by ELISpot before and after challenge (SFC – spot forming cell). Correlations between IFN-γ response and clinical scores after challenge are shown on the plots. (B) T-cell responses upon ASFV *in vitro* restimulation were analyzed by intracellular cytokine staining. (C) Correlations between the frequencies of cytokine-producing T-cell subsets and clinical scores after challenge are displayed in dot plots (r – correlation coefficient; dark red indicates a negative correlation with protection, which reflects poor clinical outcomes, while gray indicates a positive correlation with protection; correlations with p < 0.05 are outlined with black circles). In A, the data points represent values for individual animals. Measurements were made in triplicate; bars represent mean values for each animal. In B, the data points represent values for individual animals. At -9 and 7 dpc, n = 5 pigs in farm and SPF groups, and at 18 and 26 dpc, n = 2 in farm group and n = 4 in SPF group. Differences between farm and SPF groups (*), as well as between protected and non-protected animals (§) were analyzed at each time point by unpaired t-test with Holm-Šidák’s correction for multiple comparisons; *p < 0.05; **p < 0.01; ^§^p < 0.05; ^§§^p < 0.01.

Since differences in SLA (swine leukocyte antigen) alleles can shape the T-cell response by influencing peptide presentation to T cells, we studied the pigs’ haplotypes. The analysis revealed alleles found exclusively in farm or SPF pigs, as well as alleles shared between groups (Supplementary Fig. 11A). In general, the number of identified alleles per haplotype was higher in SPF pigs. However, it was not possible to associate specific haplotypes with protection, as some protected and non-protected pigs were sharing the same set of alleles (Supplementary Fig. 11B).

### Anti-p72 and hemadsorption-inhibiting antibodies are weak correlates of protection

Considering the potential role of antibodies in protection against ASFV, we measured titers of anti-p72 capsid protein antibodies using a competitive commercial ELISA. To obtain semiquantitative data, we identified the non-saturating serum concentration by serially diluting sera from a randomly selected protected SPF and a non-protected farm pig. As 50-80% reactivity was measured at 1:1250, we selected this dilution to test all sera (Supplementary Fig. 12A). The levels of anti-p72 antibodies were significantly higher in SPF pigs at all three time points, but the data did not separate protected from non-protected pigs (Supplementary Fig. 12B). Accordingly, the correlation analysis showed significant but weak correlation with protection at 169 dpi and 4 dpc (Supplementary Fig. 12C). In addition to that, we tested the levels of hemadsorption-inhibiting antibodies and again found higher titers in SPF pigs at 4 dpc, but without clear difference between protected and non-protected animals within the two groups (Supplementary Fig. 13A, B). The titers correlated significantly but weakly with protection at 4 dpc (Supplementary Fig. 13C).

### Potent activation of innate immunity and prolonged upregulation of cell cycle BTMs after Estonia 2014 infection in SPF pigs

Our follow-up investigations were driven by the hypothesis that differences in innate immunity during the immunization phase (Estonia 2014 infection) are responsible for the differences in adaptive immune responses and protection against virulent ASFV. We compared the innate responses of SPF and farm pigs using blood transcription modules (BTMs). They were computed using Gene Set Enrichment Analysis (GSEA) for ranked lists of differentially expressed genes (DEGs) of farm or SPF pigs relative to baseline (day 0). In contrast to the farm group, where induction of innate BTMs was observed mainly at 4 dpi, SPF pigs had a stronger and more prolonged activation of antigen presentation, antiviral, myeloid and NK cell BTMs lasting until 15 dpi (Fig. 4A). Regarding adaptive modules, the upregulation of cell cycle genes was more pronounced at 7 and 11 dpi in the farm group but was halted at 15 dpi. On the contrary, SPF pigs showed an extended induction of cell cycle BTMs coupled with a more robust activation of the “Signaling in T cells” along with plasma cell BTMs (Fig. 4B). This putative extended phase of lymphocyte expansion in SPF pigs was also supported by continued induction of “Respiratory electron transport chain” BTMs (Supplementary Fig. 14A). In addition to that, one of the to-be-annotated (TBA) modules (M137) was steadily induced in SPF pigs after the immunization (Supplementary Fig. 14B).

**Fig. 4.**
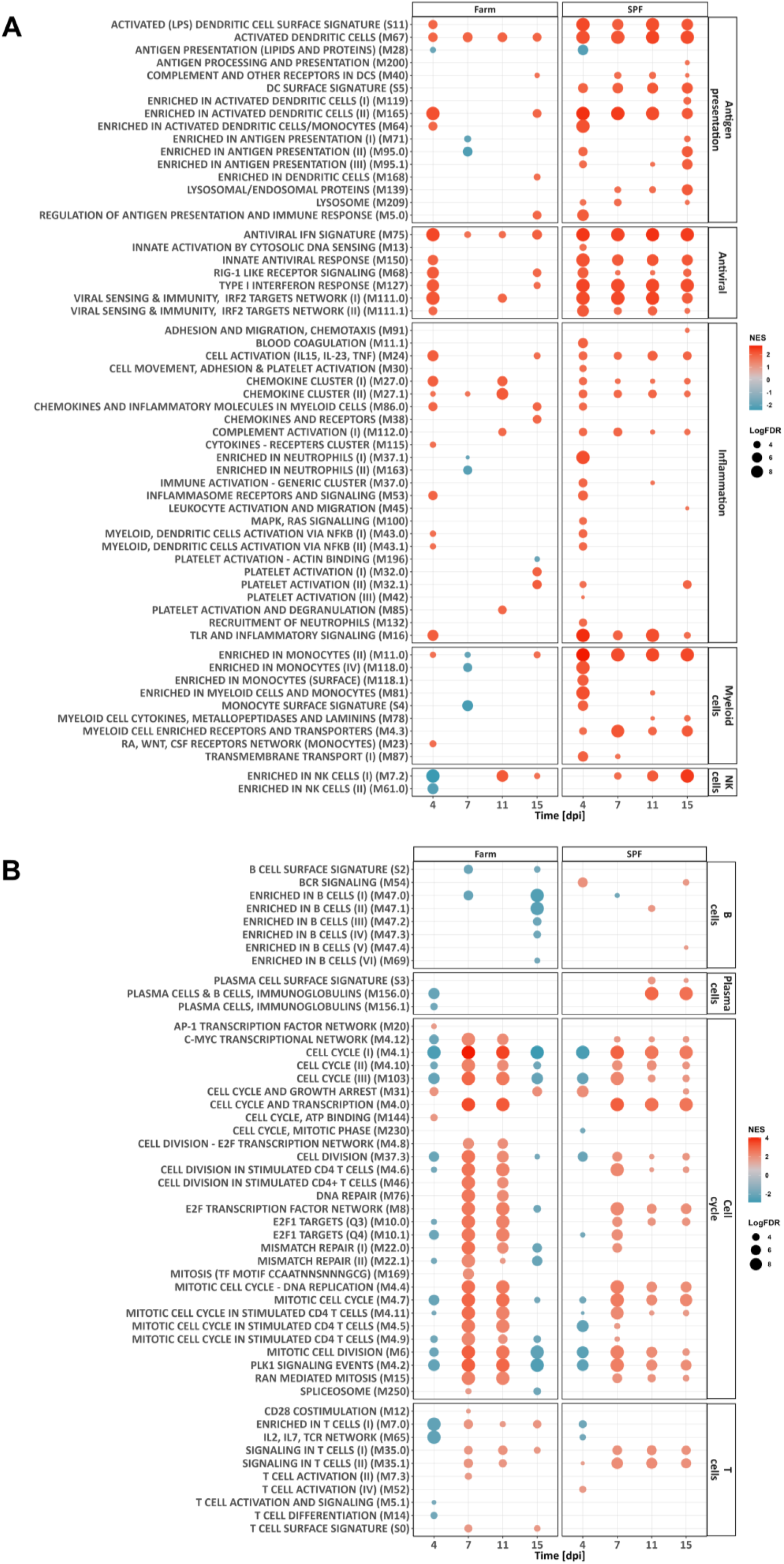
Blood leukocyte transcriptomic profiles after immunization with the Estonia 2014 strain. For each group of animals, comparisons were made against the baseline measurements (day 0). Pre-ranked list of DEGs was subjected to GSEA using BTMs as gene sets. (A, B) Dot plots show innate and adaptive BTMs, respectively. Dot size depicts the q-value (FDR), while color represents the normalized enrichment score (NES). FDR < 0.05 was selected as a cutoff.

To better understand the relationship of the blood transcriptome to events in lymphoid tissue, we also performed RNA sequencing of the gastrohepatic lymph node and spleen samples at 17 dpi. BTM scores were calculated by comparing the data of infected and mock animals of either farm or SPF origin. Upregulated antiviral BTMs were detected in both groups of pigs and in both types of organs. In contrast, only infected farm animals had a prominent activation of antigen presentation, inflammation, and myeloid cell BTMs in the lymph nodes, but not in the spleens (Supplementary Fig. 15A). In line with the PBMC data, cell cycle genes were downregulated in the lymph nodes of the farm animals, although the same modules were upregulated in the spleens of both groups of pigs. Of note, T-cell BTMs were activated only in the spleens of SPF pigs after the infection (Supplementary Fig. 15B). These results indicate that immunological events in lymph nodes and spleen are not synchronized and are strongly impacted by the hygiene status of the pigs. The transcriptomes of these organs only partially reflect the blood data.

### Sustained induction of innate immunity, plasma cell and cell cycle BTMs during the immunization phase correlates with protection

To define BTM expression patterns correlating with protection, we analyzed the relationship between clinical scores and BTM enrichment scores for individual pigs determined by Gene Set Variation Analysis (GSVA). First, enrichment scores were calculated relative to the baseline by normalizing the reads to day 0 before performing GSVA (Supplementary Fig. 16, 17). Early and durable upregulation of various antigen presentation, inflammation and myeloid cell BTMs combined with sustained induction of the antiviral M111.0 module at 11 and 15 dpi correlated with protection against challenge infection (Fig. 5A). In terms of adaptive immunity, upregulated plasma cell BTMs at 11 and 15 dpi, as well as activation of B-cell and cell cycle modules at 15 dpi, correlated with induction of protective immunity against challenge infection (Fig. 5B).

**Fig. 5.**
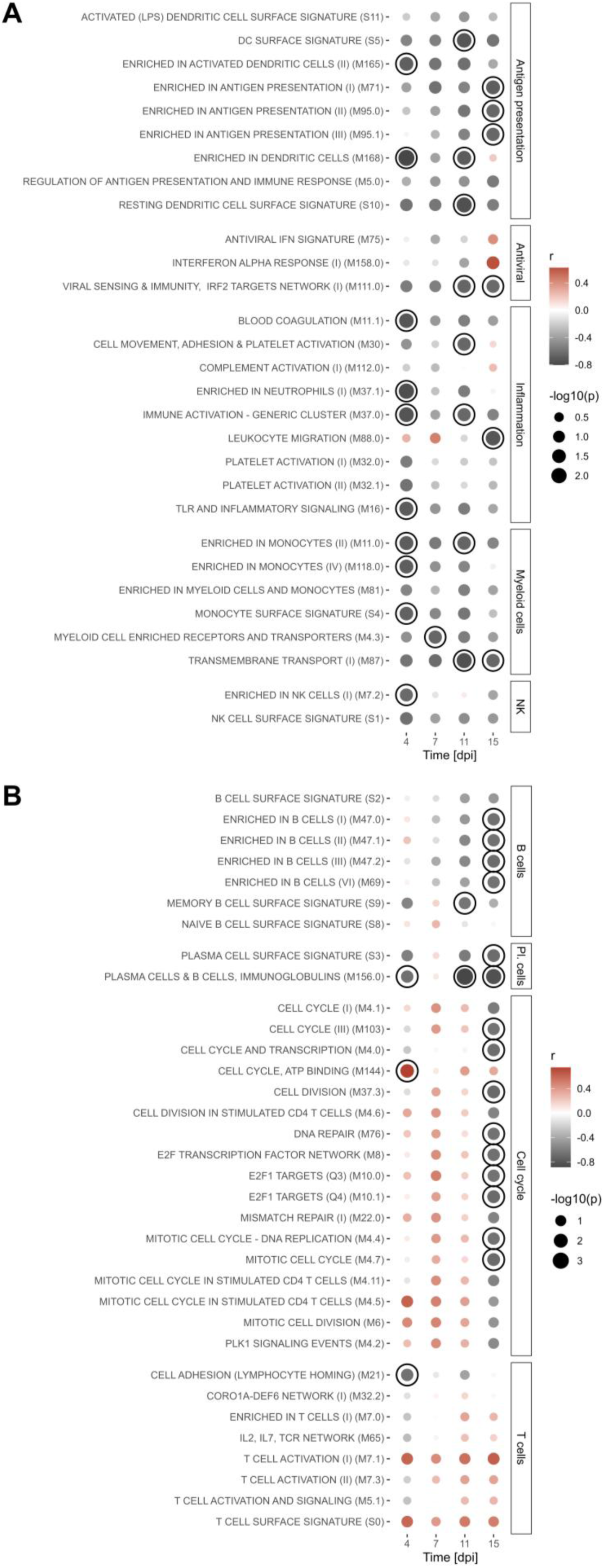
Correlation of relative BTM expression after Estonia 2014 immunization with protection. Read counts after bulk RNA-Seq were normalized by the baseline values (day 0). GSVA was then performed to calculate NES values for individual animals using BTMs as gene sets. Obtained NES values were correlated with AUC values of the challenge clinical outcomes. In A and B, the dot plots show innate and adaptive BTMs, respectively. Dot size depicts the p-value, and color represents the correlation coefficient r (dark red indicates a negative correlation with protection, which reflects poor clinical outcomes, while gray indicates a positive correlation with protection). Correlations with p < 0.05 are outlined with black circles.

### Higher baseline levels of innate immunity BTMs correlate negatively with protection, while expression of adaptive immunity modules has the opposite effect

While Fig. 5 represents correlation of relative BTM values with protection, we also performed similar analysis with absolute enrichment scores of BTMs computed by GSVA using non-normalized (absolute) read counts shown on heatmaps in Supplementary Fig. 18-20. The calculations demonstrated that baseline expression of antigen presentation, antiviral, inflammation, and myeloid cell modules correlated negatively with protection against challenge infection, which can be explained by the higher expression levels of these BTMs in farm pigs at baseline (Supplementary Fig. 18). Nevertheless, it was remarkable to see that expression of several innate BTMs correlated with worse clinical outcomes up until 15 dpi. Amongst the innate BTMs, only the induction of the antigen presentation module M168 at 4 dpi and the NK cell module M7.2 at 4 and 15 dpi correlated with protection (Fig. 6A).

**Fig. 6.**
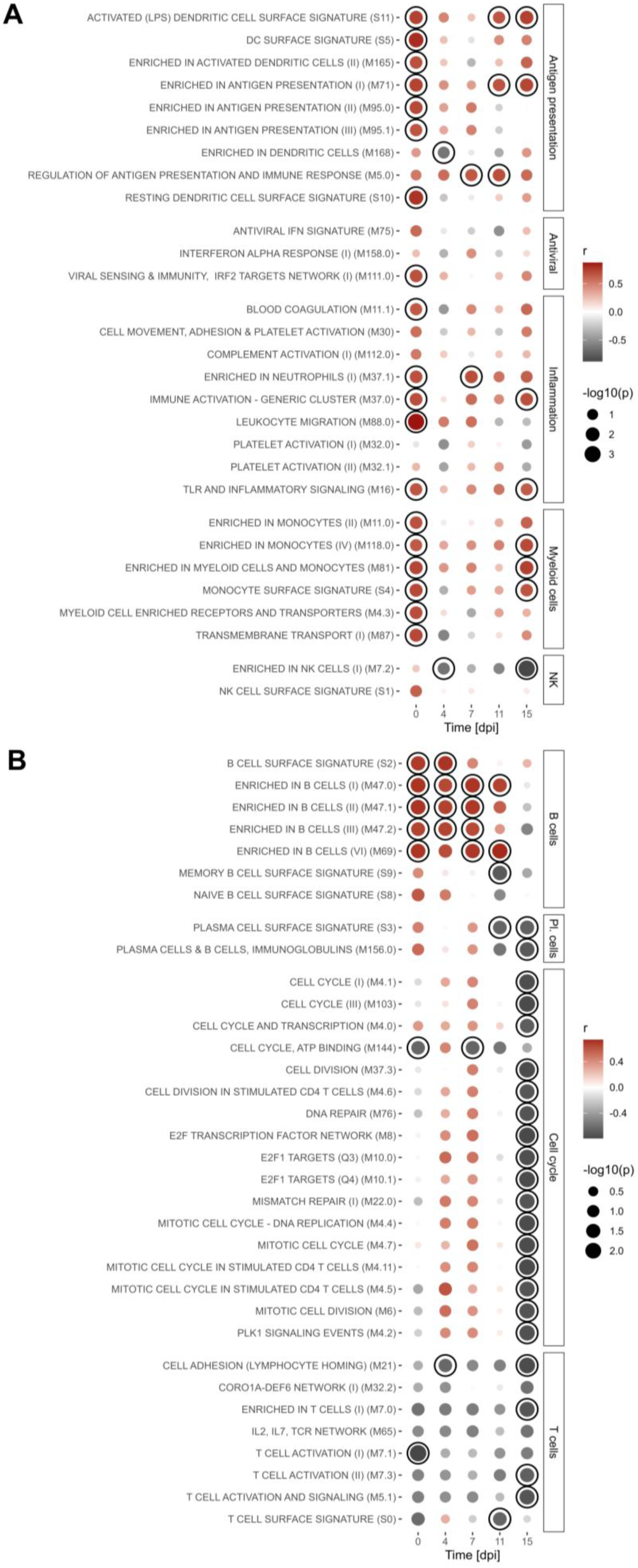
Correlation of absolute BTM expression after Estonia 2014 immunization with protection. GSVA was performed on non-normalized read counts to calculate NES values for individual animals using BTMs as gene sets. Obtained NES values were correlated with AUC values of the challenge clinical outcomes. In A and B, the dot plots show innate and adaptive BTMs, respectively. Dot size depicts the p-value, and color represents the correlation coefficient r (dark red indicates a negative correlation with protection, which reflects poor clinical outcomes, while gray indicates a positive correlation with protection). Correlations with p < 0.05 are outlined with black circles.

Regarding adaptive BTMs, we found negative associations with protection at 0-11 dpi for absolute B-cell module scores. This contrasted with the expression of plasma cell BTMs at 11 and 15 dpi that correlated with better clinical outcomes. In addition, absolute scores of cell cycle and several T-cell BTMs at 15 dpi represented correlates of protection. Notably, correlations with disease resistance, although often not significant, were observed for most T-cell modules across all time points (Fig. 6B).

To better understand the apparent discrepancies between correlations using GSVA scores calculated relative to the baseline versus absolute GSVA scores, we plotted the absolute scores of 10 animals with known clinical outcomes across 5 time points, grouping them into a single mean value for each BTM family (Fig. 7). In line with entire group comparisons (Supplementary Fig. 18), SPF animals had lower baseline levels of antigen presentation, antiviral, inflammation, and myeloid cell BTMs. However, because of a more potent upregulation of these innate modules, their expression rates became comparable to the values of the farm group at 4 dpi. NK and T-cell modules were steadily induced from 4 to 15 dpi, with no clear separation between protected and non-protected animals in either group. Similarly, the expression of B cell-related genes did not distinguish between protected and non-protected animals in the two groups; instead, it reflected baseline differences, which diminished by 11 dpi. Strikingly, expression patterns of plasma cell modules contrasted in protected versus non-protected farm animals, being significantly higher in protected pigs at 11 dpi. The enhancement of these modules was also observed in SPF pigs at 11 and 15 dpi that matched with prolonged activation of cell cycle genes in this group.

**Fig. 7.**
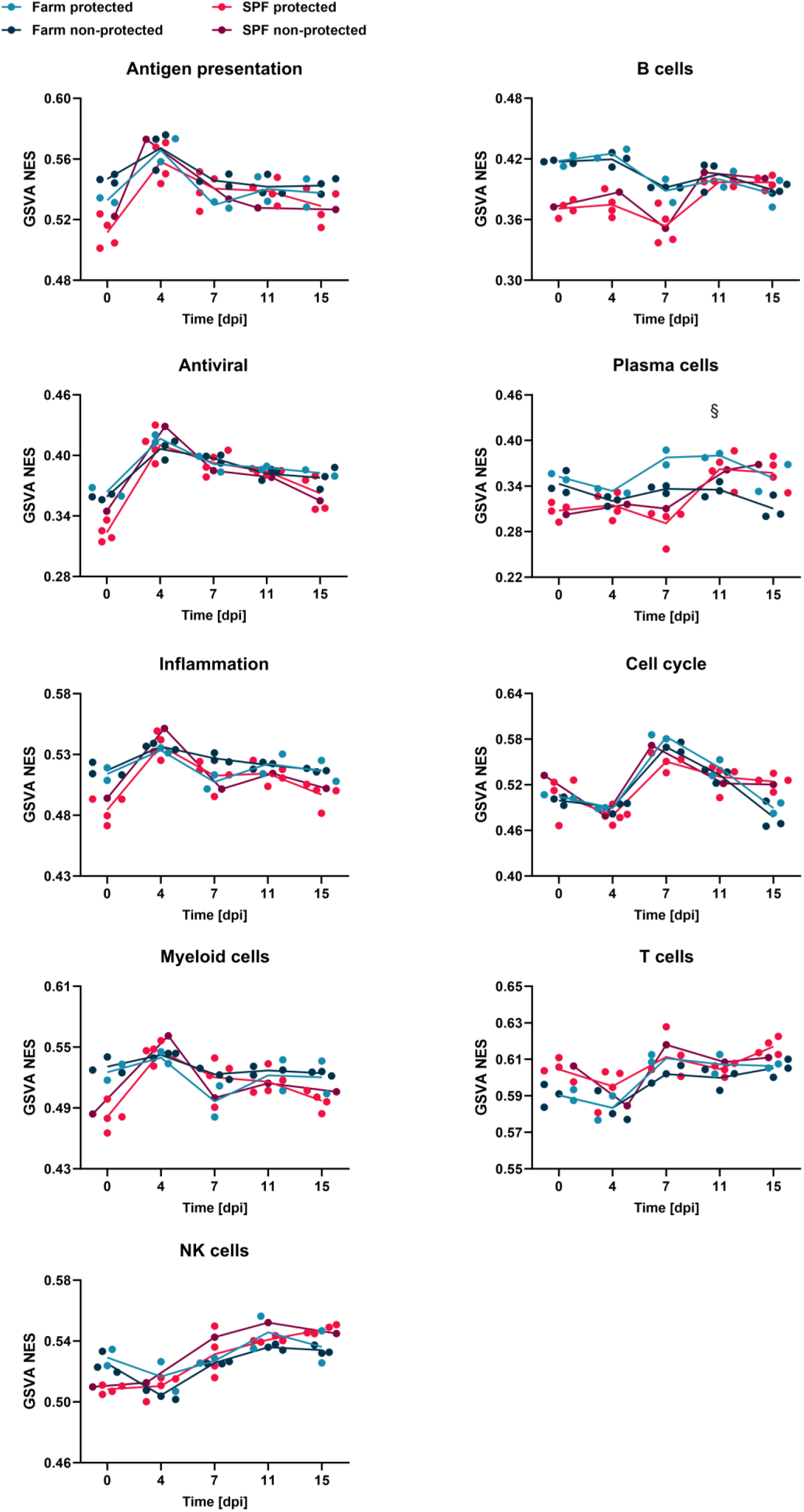
Kinetics of average NES of BTM families after Estonia 2014 immunization. NES values were calculated by GSVA for five farm and five SPF pigs using non-normalized read counts. At each time point, NES values of BTMs belonging to one family were used to determine average family scores. Plots demonstrate average NES values for each family related to either innate (left column) or adaptive immunity (right column). Both farm and SPF groups had n = 5 pigs each. Differences between protected and non-protected farm pigs were analyzed by unpaired t-test with Holm-Šídák’s correction for multiple comparisons; ^§^p < 0.05.

### PBMC transcriptome alterations after challenge indicate that continued effector T cell and plasma cell generation correlates with protection

Next, we correlated post-challenge BTM scores calculated by GSVA using absolute read counts (Supplementary Fig. 21, 22) with the clinical outcomes. The induction of innate immunity BTMs at 7 dpc generally negatively correlated with protection, although the upregulation of the “Platelet activation (I) (M32.0)” module at 11 dpc was linked to better clinical outcomes (Fig. 8A). As for adaptive BTMs, we observed a trend of positive correlation between early induction of cell cycle modules (4 dpc) and disease resistance. Activated plasma cell and T-cell modules between 4 and 11 dpc represented the main correlates of protection. In contrast, delayed upregulation of cell cycle and some of the B-cell modules at 7 dpc negatively correlated with protection (Fig. 8B). The temporal expression of innate and adaptive BTMs for individual animals indicated that the non-protected SPF pig #1967 differed from the protected pigs in terms of immune response. It had a particularly strong induction of innate BTMs at 7 dpc that appeared to be associated with a drop in adaptive immunity activation not seen in any other pig (Supplementary Fig. 23).

**Fig. 8.**
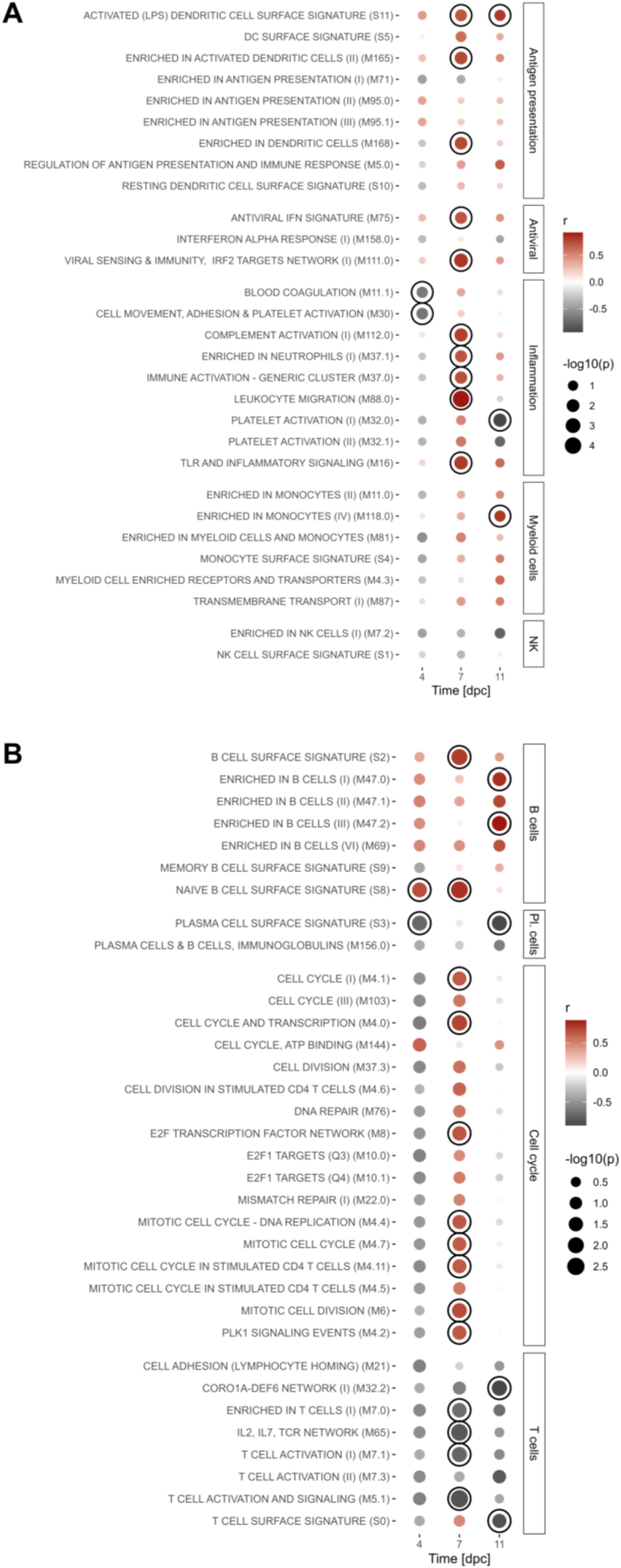
Correlation of absolute BTM expression after Armenia 2008 challenge with protection. GSVA was performed on non-normalized read counts to calculate NES values for individual animals using BTMs as gene sets. Obtained NES values were correlated with AUC values of the challenge clinical outcomes. In A and B, the dot plots show innate and adaptive BTMs, respectively. Dot size depicts the p-value, and color represents the correlation coefficient r (dark red indicates a negative correlation with protection, which reflects poor clinical outcomes, while gray indicates a positive correlation with protection). Correlations with p < 0.05 are outlined with black circles.

## Discussion

At present, the development of an effective and safe vaccine against ASFV is of utmost importance. LAVs have demonstrated promising efficacy in recent studies (Urbano and Ferreira, 2022; Vu and McVey, 2024); however, the balance of the immune response between protection and immunopathology remains a critical issue (Chu et al., 2024). For that reason, it is essential to understand protective and detrimental immune responses following vaccination and challenge infection. In the present study, we employed a systems immunology approach to generate a temporally resolved model of protective innate and adaptive immune responses based on multiple correlation analyses (Fig. 9). We used age-, sex-, and breed-matched pigs that were either of SPF or non-SPF (farm) origin. The model is based on our previous data regarding the impact of baseline immune status on ASF pathogenesis and the development of protection (Radulovic et al., 2022, 2025). In the latter and present studies, SPF pigs developed more robust protective immunity compared to farm animals following immunization with the ASFV Estonia 2014 strain.

**Fig. 9.**
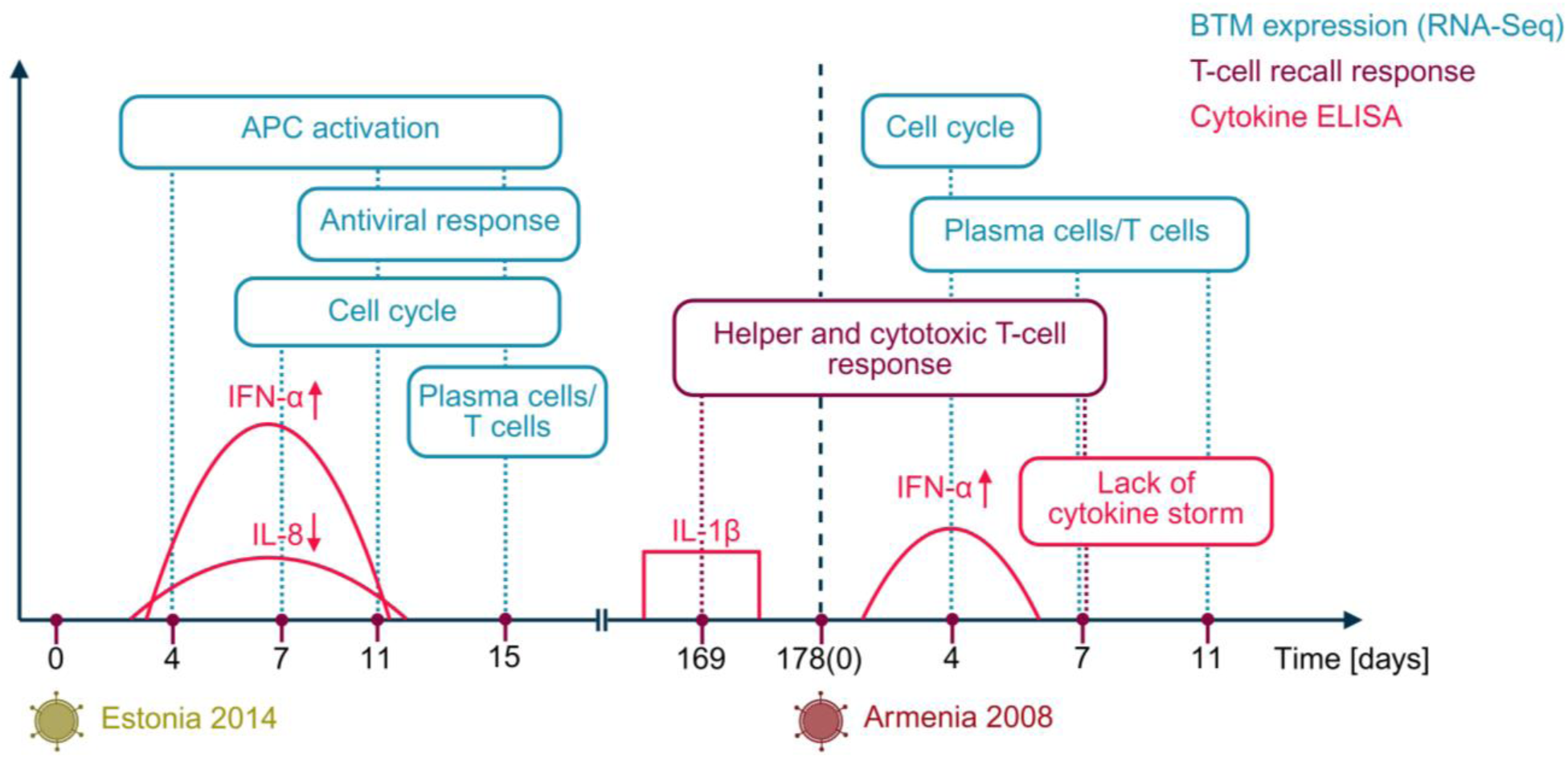
Model of protective immune responses against ASFV challenge in immunized pigs. The findings from BTM induction, systemic cytokine, and T-cell response analyses are integrated. Following immunization with the attenuated virus, an early IFN-α response activates antigen-presenting cells (APCs), triggering sustained cell cycle activation and clonal expansion of antigen-specific lymphocytes. This process generates plasma cells and effector T cells, with efficient memory T helper cell development emerging as a key correlate of protection. Upon challenge, a controlled IFN-α response and a rapid activation of cell cycle genes correlate with protection. The coordinated action of antibody-producing plasma cells and activated helper and cytotoxic T cells effectively controls viral replication while preventing a cytokine storm, ultimately conferring protection against ASFV.

Our data suggest an important role for the early type I IFN response during both immunization and challenge phases. The more prominent IFN-α release in protected pigs after the immunization with the Estonia 2014 strain did not reduce virus replication and spread sufficiently to induce differences in viremia and viral loads in organs, indicating that the cytokine acted more via alarming innate immunity to promote antigen-presenting cell (APC) activation, and thereby adaptive immune response induction (Crouse et al., 2015; Dahlgren et al., 2022). This is supported by the fact that prolonged activation of antiviral BTMs correlated with protection and was associated with upregulation of antigen presentation and adaptive immunity modules, as well as enhanced T-cell and antibody responses. Following challenge infection, only the early IFN-α response (4 dpc) correlated positively with protection, while antiviral BTMs at 7 dpc correlated negatively. This is understandable since non-protected pigs did not control virus replication, which likely sustained IFN-α production. Based on previous works by our group and others, we propose that plasmacytoid dendritic cells (pDCs) are the source of IFN-α in the context of ASFV infection, in particular considering that IFN-α responses are efficiently suppressed in virus-infected macrophages (Golding et al., 2016; Sánchez-Carvajal et al., 2025). Further identification of IFN-α-producing cells in diverse tissues following ASFV infection is required to confirm this *in vivo,* as it has been demonstrated for classical swine fever virus (Auray et al., 2020). In addition, characterization of pDC interaction networks will provide valuable information to better understand antiviral immunity mechanisms (Van Eyndhoven et al., 2021).

Previous exposure to pathogens is likely responsible for a pro-inflammatory baseline status of farm pigs when compared to SPF pigs, and our data indicate that this difference impacts the response to ASFV infection. Farm pigs had higher baseline levels of pro-inflammatory IL-1β and IL-8, coupled with enhanced innate immunity BTM expression that represented a negative correlate of protection. To understand how inflammatory responses may impact protection or disease progression, both absolute (calculated at each time point) and relative (normalized to day 0 post-immunization) BTM enrichment scores were considered. Although SPF pigs had a lower baseline innate immunity status, the upregulation of the corresponding BTMs early after the immunization was more potent than in the farm group, which represented a robust correlate of protection. This may stem from the differences in leukocyte functionality and distribution in tissues between farm and SPF animals that can impact in varying degrees the quality and quantity of their immune responses (Beura et al., 2016; Magden et al., 2020). According to existing knowledge, a higher pro-inflammatory status during the immunization phase could lead to impaired T and B cell differentiation and function (Danahy et al., 2016; Duan et al., 2020). Indeed, excessive production of IL-1β can cause dysregulation of adaptive immunity, and IL-8 promotes inflammation through neutrophil recruitment and activation (Xiang et al., 2023).

The role of cytokines can vary depending on the time during the immunization-challenge experiment. For example, a moderate level of IL-1β before challenge (169 dpi) was shown to be a correlate of protection. The function of this cytokine is potentially different at this time point compared to the period after immunization with the ASFV Estonia 2014 strain. It has been previously shown that IL-1β acts as a licensing signal for memory CD4^+^ T cell function (Jain et al., 2018). Indeed, non-protected farm pigs with the lowest levels of IL-1β before the challenge had almost undetectable cytokine responses from SP and DP T_h_ cells. Defining the conditions and cellular sources of IL-1β secretion at this stage of infection should therefore be addressed in future studies. On the one hand, it may result from low-level viral persistence in tissues. On the other hand, it could reflect a residual, primed immune state following immunization. In the latter case, macrophages with a trained phenotype could secrete IL-1β and display enhanced responsiveness upon challenge (Taks et al., 2022; Bahl et al., 2025). Conversely, increased levels of IL-1β at 7 dpc, as well as IL-1RA, IL-8, and IFN-γ at 7 and 11 dpc, negatively correlated with protection. The detrimental pro-inflammatory response was also reflected in the blood transcriptome data: the majority of BTMs associated with inflammation negatively correlated with protection at 7 dpc. It is worth noting that expression of the module “Platelet activation (I) (M32.0)” at 11 dpc correlated with enhanced resilience to challenge infection. A potential role of platelets in modulating antibody response durability was recently highlighted by Cortese et al. (2025), yet the exact role of platelets in protection against ASFV remains to be elucidated.

NK cell activity after ASFV infection was shown to depend on the strain’s virulence, with an increase in their cytotoxic functions mostly observed against non-pathogenic strains (Takamatsu et al., 2013; Norley and Wardley, 1983; Leitão et al., 2001). Our blood transcriptome data demonstrated that upregulation of the module “Enriched in NK cells (I) (M7.2)” after the immunization phase positively correlated with protection. In addition, increased NK cell activation in terms of IFN-γ production was observed post-challenge in protected farm animals (Supplementary Fig. 8). Therefore, the role of NK cells in protection against genotype II ASFV should be explored in future studies.

Our study demonstrates that T cell activation upon immunization and challenge represents a robust correlate of protection against ASFV. The expression of T-cell BTMs between 0 and 15 dpi positively correlated with improved clinical outcomes, most prominently at 15 dpi. Along with that, upregulation of cell cycle modules at 15 dpi, possibly reflecting a clonal expansion of lymphocytes, was also shown to correlate with protection. Nonetheless, the abrogation of cell cycle BTMs in blood at 15 dpi and in lymph nodes and spleen at 17 dpi was a prominent feature in farm pigs.

It is important to note that the IFN-γ response from freshly collected PBMCs detected by ELISpot prior to challenge did not reflect the outcome when correlation analysis was performed across both farm and SPF groups. Still, protected farm pigs had elevated cytokine secretion when compared to non-protected ones within the same group. Flow cytometric analysis of PBMC restimulation with ASFV revealed differences in IFN-γ and TNF production prior to challenge and enabled a more in-depth characterization of cellular responses. While the increase in SP and DP T_h_, along with T_c_ cell populations, after ASFV infection was shown previously (Schäfer et al., 2021), we demonstrate that effector cytokine responses from these cell types have distinct patterns and strongly correlate with protection. A more potent TNF production by DP T_h_ cells was observed in the SPF group before the challenge, which might have potentially promoted antibody production by plasma cells. Alternatively, TNF was shown to induce metabolism-dependent antiviral state in target cells, which could also be applied in the context of ASFV (Ciesla et al., 2022). DP are memory CD4^+^ T cells that start to express CD8α after differentiation. Despite their importance in protection against ASFV, their exact functions remain largely unknown (Schäfer et al., 2022). The protective activation of T_c_ cells was, however, more pronounced at 7 dpc, likely marking the elimination of virus-infected cells. The cytokine response from effector γδ T cells was a weaker correlate of protection, although IFN-γ production at 7 dpc was detected only in the survived animals. Besides secretion of cytokines and chemokines, these cells may be involved in antigen presentation (Takamatsu et al., 2006) and may potentially mediate cytotoxicity, as they have been shown to express perforin after ASFV infection (Hühr et al., 2020).

Since T cells can also mediate immunopathology, an important finding of our study was that ASFV-specific effector T-cell responses, combined with T-cell BTM expression, correlated with protection at 7 dpc. The positive correlation for BTMs at 4 dpc followed the same trend, however, it was not significant enough. This was due to a strong but only temporary induction of T-cell modules in the non-protected SPF pig #1967. However, unlike T cells, prolonged induction of cell cycle BTMs at 7 dpc correlated with worse clinical outcomes, indicating a possible delay in clonal lymphocyte expansion. It was pointed out in a previous study by Sánchez-Cordón et al. (2020) that the absence of long-term protection following immunization with attenuated genotype I ASFV strains may be linked to the induction of immunosuppression, characterized by increased numbers of regulatory T cells and elevated IL-10 levels in serum. Unlike immunization with vaccine strains, Estonia 2014 infection causes prolonged viremia in pigs, which might contribute to T cell exhaustion. Nevertheless, while lesions were observed in organs collected at 17 dpi in the present study, no pathological signs were detected in organs collected at 28 dpi in the previous study by Radulovic et al. (2022). To definitively rule out the role of immunosuppression, markers of T cell exhaustion should be included in immunophenotyping for subsequent studies.

Besides the role of T cells, it is important to understand the contribution of antibodies to protection against ASFV. Indeed, antibody-mediated responses represented correlates of protection in our study. The expression of plasma cell BTMs at 11 and 15 dpi, as well as at 4 and 11 dpc, correlated with improved clinical outcomes. In addition, titers of hemadsorption-inhibiting antibodies quantified at 4 dpc and anti-p72 antibodies quantified pre-challenge and at 4 dpc positively correlated with protection as well. It was previously demonstrated that ASFV virions can exist without the outer envelope and remain infectious (Gao et al., 2023). Accordingly, when exposed to anti-capsid p72 antibodies, such virions could be potentially neutralized. Of course, our data do not allow any conclusion on the function of anti-p72 antibodies in protection, and the titer increase may only reflect enhanced T_h_ cell response. Furthermore, since ASFV is only partially neutralized by antibodies (Escribano et al., 2013), alternative mechanisms such as ADCC, ADCP, CDC, or intracellular infection inhibition should be further investigated, particularly for genotype II strains (Wardley and Wilkinson, 1985; Foss et al., 2019; Chen et al., 2025). In this context, hemadsorption-inhibiting antibodies target viral surface proteins CD2v and C-type lectin and thus can be immunologically relevant (Malogolovkin et al., 2015; Burmakina et al., 2016; Gladue et al., 2020). It would be worth studying them in more detail, despite the poor correlation with protection observed that might be related to the assay used. Certainly, our data support future experiments, such as passive transfer of immune serum, to elucidate the role of antibodies in protection, as recently performed in a study by Friedrichs et al. (2025a) for genotype II strains. Besides that, more immunogenic ASFV proteins and functions of protective antibodies targeting them need to be identified.

Generally, the low percentage of protected farm pigs can be partially attributed to the short duration of immunity, as the challenge time point in the present study was selected relatively late, based on the clearance of the virus from blood and serum. However, in a study by Friedrichs et al. (2025b, in revision), farm pigs immunized with the Estonia 2014 strain and challenged with the Armenia 2008 strain after six months demonstrated a high level of protective immunity. It would likely be informative to also examine the early events following immunization with the Estonia 2014 strain or a LAV candidate, by assessing the onset of immunity (Marín-Moraleda et al., 2024) or more thoroughly characterizing the kinetics of adaptive immune responses, as well as the modulation of immune memory formation by viral persistence. Notably, based on our unpublished data, persistence of the LAV strain did not reduce protection against ASFV challenge in farm pigs.

In summary, our data provide a framework of protective immune responses following immunization with the attenuated ASFV Estonia 2014 strain and subsequent challenge with the pathogenic Armenia 2008 strain, both belonging to genotype II (Fig. 9). Based on the identified correlates, we propose that a relatively sustained IFN-α response promotes APC activation ensuring sufficient clonal expansion of specific lymphocytes during at least two weeks. Consequently, the induction of both plasma cell and T-cell BTMs at 15 dpi correlated with protection. As expected from this pattern, the activation of virus-specific memory T cells, in particular T_h_ cells, and anti-ASFV antibody responses detected at the time of challenge, represented correlates of protection. Of note, further activation of T cells after the challenge infection positively correlated with better clinical outcomes indicating that T cells do not cause immunopathology during ASFV and should be targeted in vaccine design.

Our experimental model has several limitations that need to be addressed in future studies. While SPF animals provide a controlled environment for studying specific immune responses, they may not reflect how the immune system functions in the field conditions (Chen et al., 2023). In a farm setting, pigs may have varying levels of innate immune activation and different profiles of disease exposure, both of which can influence the immune response to a vaccine or infection (Salines et al., 2019; Chrun et al., 2023). Notably, in the previous study on protective immunity against ASFV in SPF versus farm pigs, disease severity after Estonia 2014 oronasal infection was significantly lower in SPF than in farm animals (Radulovic et al., 2025). This contrasts with the course of Estonia 2014 infection observed in the present study. The farm pigs used in the previous study came from a different source than those used here. Therefore, the identified correlates should be verified using pigs originating from a broad range of farm environments. It should also be noted that the ASFV Estonia 2014 strain does not represent a vaccine strain and still retains considerable virulence. For this reason, the correlates should also be validated with vaccine candidates. Independent of these open questions, the present work demonstrated a strong impact of the immunological baseline on development of antiviral immunity. The observation that a more quiescent immunological status may be advantageous is surprising, especially considering the concept of trained immunity, which supports the beneficial effects of prior innate immune activation in protection against pathogens (Ziogas et al., 2023).

## Methods

### Ethics Statement

The experimental infection of the pigs was performed according to the Animal Welfare Act (TSchG SR 455), the Animal Welfare Ordinance (TSchV SR 455.1), and the Animal Experimentation Ordinance (TVV SR 455.163) of Switzerland. The animal experiment was reviewed by the committee on animal experiments of the canton of Bern and approved by the cantonal veterinary authority under the licenses BE18/2019 and BE46/2022.

### Animal experiment

A total of 24 male and female Large White domestic pigs (age, 10-11 weeks; weight, ∼20-25 kg) were obtained from the IVI SPF facility (n = 12; termed “SPF pigs”) and from Agroscope’s Posieux experimental farm (n = 12; termed “farm pigs”). The SPF pigs were not vaccinated and were negative for all porcine viral and bacterial pathogens. The farm pigs were vaccinated against porcine circovirus type 2, porcine parvovirus K22, *Escherichia coli*, and *Erysipelothrix rhusiopathiae*. The animals were moved to the BSL3-Ag containment facility of the IVI and randomly assigned to virus-infected (n = 18) or mock (n = 6) groups, with SPF and farm pigs kept in separate stables. After 5 days of acclimatization, the pigs were oronasally infected with 5 mL of blood containing 2.51×10^7^ 50% tissue culture infectious dose (TCID_50_) of the attenuated ASFV Estonia 2014 strain per animal. The mock animals received 5 mL of non-infected blood. Three pigs in each group were sacrificed at 17 days post-immunization (dpi). The mock animals were euthanized at 30 dpi. One immunized animal in the farm group was euthanized at 74 dpi due to a bacterial infection, while one in the SPF group was euthanized at 149 dpi to free up space in the stable. The remaining five pigs in each group were oronasally challenged with the highly virulent ASFV Armenia 2008 strain (3.97×10^7^ TCID_50_ in 5 mL blood per animal) at 178 dpi, which corresponded to day 0 post-challenge (dpc). At the end of the experiment (26 dpc or 204 dpi) or when the criteria for discontinuation were reached (see below), the pigs were euthanized by electrical stunning and complete blood withdrawal by a professional butcher certified for stunning and exsanguination of livestock. Rectal temperatures and clinical scores were recorded daily by a veterinarian. Blood samples were taken for viremia monitoring, measurement of cytokines and ASFV-specific antibodies, leukocyte isolation, and bulk RNA extraction. Organs were collected for pathological examination, viral load quantification, and bulk RNA extraction.

### Clinical scoring

A well-established clinical scoring system described by Mittelholzer et al. (2000) was employed to assess pig health and welfare, evaluating nine parameters on a scale from 0 (normal) to 3 (severe abnormality). The assessment included: Liveliness (0: attentive, curious, stands up immediately; 1: slightly reduced, stands up hesitantly; 2: tired, gets up only when forced; 3: somnolence for 24 h or body temperature <37.6°C); Body tension/posture (0: relaxed, straight back; 1: stiffness and bent back while standing up, afterwards normal; 2: bent back and stiff walking remains; 3: seizures or cramps of neurological origin, unable to walk); Body shape (0: full stomach, "round" body; 1: empty stomach; 2: empty stomach, thinned body muscles; 3: emaciated, backbone and ribs clearly visible); Respiration (0: frequency 10-20/min, barely visible chest movement; 1: frequency >20/min; 2: frequency >30/min, distinct chest movement; 3: frequency >40/min and breathing through open mouth); Walking/gait (0: well-coordinated movements; 1: hesitant walking, crossed-over legs corrected slowly; 2: distinct ataxia/hind lameness, able to walk; 3: paresis/paralysis, unable to walk for 24 h); Skin (0: evenly light pink, hair coat flat; 1: reddened skin areas; 2: clearly discolored cold skin areas, petechial hemorrhages; 3: purple or black-red discoloration, no sensitivity, large hemorrhages in skin and/or nosebleed); Eyes/conjunctiva (0: light pink; 1: light conjunctivitis, clear secretion; 2: moderate conjunctivitis, turbid secretion; 3: severe conjunctivitis, purulent secretion); Appetite (0: greedy, hungry; 1: eats slowly when fed; 2: doesn’t eat when fed, but sniffs food; 3: doesn’t eat at all, shows no interest for food); and Defecation (0: soft feces, normal amount; 1: reduced amount of feces, dry; 2: only small amount of dry, fibrin-covered feces, or diarrhea; 3: no feces, mucus in rectum, or watery and bloody diarrhea).

### Virus stocks and quantification

The genotype II ASFV strains Estonia 2014 (Zani et al., 2018) and Armenia 2008 (Gabriel et al., 2011) were kindly provided by Sandra Blome and Martin Beer (Friedrich-Loeffler-Institut, Greifswald-Insel Riems, Germany). Virus titers were determined by immunofluorescence assay in WSL cells (Keil et al., 2014) as previously described by Radulovic et al. (2022). The ASFV DNA was isolated from samples using the NucleoMag VET kit (Macherey-Nagel, Germany) and a Kingfisher Flex extraction robot (Thermo Fisher Scientific, USA) according to the manufacturer’s instructions. DNA extractions were performed from 200 μl of either EDTA blood, serum or organ homogenates in RA1 lysis buffer (Macherey-Nagel, Germany). Next, qPCR was performed according to the published protocol (King et al., 2003), followed by quantification of genome equivalents (Radulovic et al., 2022). Standards were analyzed in triplicate, and samples in duplicate.

### Cytokine and p72 ELISAs

The following serum cytokines were measured by conventional ELISA: IL-1α, IL-1β (RayBiotech, USA), IL-1ra, IL-2, IL-8, TNF (R&D Systems, USA), IFN-γ (Mabtech, Sweden) and IFN-α (in-house assay using the monoclonal antibodies K9 and F17 that were kindly provided by Dr. B. Charley, INRA, Jouy-en-Josas, France) (Guzylack-Piriou et al., 2006). Antibodies against the viral capsid protein p72 were detected with INgezim PPA Compac blocking ELISA (Ingenasa, Spain). The results are shown in % of inhibition with following cut-off values: ≤40% - negative; 41-49% - questionable; ≥50% - positive. Serum samples were analyzed in duplicate.

### Hemadsorption inhibition assay (HADIA)

The method is based on a published protocol with slight modifications (Malogolovkin and Sereda, 2022). IPKM cells (Masujin et al., 2021) were seeded in 96-well plates at 4 x 10^4^ cells/100 μl medium/well, infected with ASFV Georgia 2007 at MOI = 0.1, and incubated overnight at 37°C with 5% CO_2_. In separate U-shaped 96-well plates, two-fold dilutions (1:10 to 1:1280) of heat-inactivated sera in complete macrophage medium were prepared. Further, 100 μl of pre-diluted sera were transferred to the cells in six replicates and the plates were incubated for 2 h. To each well, 50 μl of 0.02% swine red blood cells were added and the plates were left overnight at 37°C with 5% CO_2_. The photos of the wells were taken by ImmunoSpot S5 UV analyzer (Cellular Technology LTD, USA). The RBC rosettes were counted using ImageJ software and the ND_50_ titers of hemadsorption-inhibiting antibodies were calculated with the Spearman-Kärber formula (Ramakrishnan, 2016).

### IFN-γ ELISpot

Freshly collected peripheral blood mononuclear cells (PBMCs) were isolated using Ficoll-Paque 1.077 g/L (Cytiva, USA) density centrifugation and used to quantify cytokine production in response to the live virus. Briefly, 96-well MultiScreen PVDF plates (Merck, Germany) were coated with a capture anti-pig IFN-γ antibody (559961, BD Pharmingen, USA) at 0.83 μg/mL in sterile PBS, incubated overnight at 4°C and blocked for 1 h at 37°C with 0.5% BSA diluted in PBS. PBMCs were counted and plated in ELISpot plates at 5 x 10^5^ cells/well (triplicate per condition) in AIM-V medium (Gibco, USA). Cells were restimulated either with the ASFV Estonia 2014 strain (MOI = 0.1) or with supernatant of mock-treated IPKM cells. After 36 h incubation at 37°C with 5% CO_2_, the plates were stained with a biotinylated detection anti-pig IFN-γ antibody (559958, BD Pharmingen, USA), streptavidin-HRP (Dako, USA) and TMB substrate (Mabtech, Sweden). The plates were scanned and analyzed using ImmunoSpot S5 UV analyzer (Cellular Technology LTD, USA).

### Intracellular cytokine staining

To analyze production of IFN-γ and TNF by several immune cell subsets, freshly isolated PBMCs were seeded in U-shaped 96-well plates at 5 x 10^5^ cells/well in AIM-V medium (Gibco, USA) and restimulated with the ASFV Estonia 2014 strain (MOI = 0.1) or mock treated for 18 h at 37°C with 5% CO_2_. Brefeldin A (eBioscience, USA) was added 4 h before cell harvesting at 1:1000 dilution. The antibody staining panel was based on a previously described protocol (Matthijs et al., 2019). Antibodies and secondary reagents are listed in Supplementary Table 1. Free binding sites of secondary antibodies were blocked with whole anti-mouse IgGs (Jackson ImmunoResearch, UK). Live/Dead Aqua stain kit (Invitrogen, USA) was used to discriminate and exclude dead cells from the analysis. Cytofix/Cytoperm kit (BD Biosciences, USA) for fixation and permeabilization was used according to the manufacturer’s instructions.

### Flow cytometry and hematology analysis

The samples were acquired on FACSCanto II flow cytometer (BD Biosciences, USA) equipped with three lasers (405, 488, and 633 nm). The data were analyzed using FlowJo v10 (FlowJo Software, USA) following the gating strategy shown in Supplementary Fig. 7. Differential blood cell counts from EDTA blood samples were measured on Vetscan HM5 hematology analyzer (Zoetis, UK).

### RNA sequencing

EDTA blood samples were first treated with ammonium chloride buffer to lyse erythrocytes as previously described by Ardali et al. (2024). Single-cell suspensions were obtained from gastrohepatic lymph nodes and spleens (Auray et al., 2020). RNA was extracted from white blood cells and from single-cell suspensions of organs using Trizol and NucleoSpin RNA kit (Macherey-Nagel, Germany). Libraries were prepared with TruSeq Stranded mRNA kits (Illumina, USA). RNA quality was controlled with a Fragment Analyzer (5200 Fragment Analyzer CE instrument, Agilent, USA). Samples were sequenced using paired-end reads of 150 bp on the Next Generation Sequencing (NGS) platform at the University of Bern using the NovaSeq 6000 sequencer (Illumina, USA). The quality of the RNA-Seq data was assessed using fastqc v.0.11.9 (Andrews, 2022) and RSeQC v.4.0.0 (Wang et al., 2012). The reads were mapped to the reference genome using HiSat2 v.2.2.1 (Kim et al., 2015). FeatureCounts v.2.0.1 (Liao et al., 2014) was used to count the number of reads overlapping with each gene, as specified by the *Sus scrofa* genome assembly (Sscrofa11.1.108). The Bioconductor package DESeq2 v1.38.3 (Love et al., 2014) was used to test for differential gene expression between the experimental groups.

### Blood transcription module (BTM) analyses

BTM analyses were performed as previously described (Matthijs et al., 2019). Briefly, after ranking of DEGs based on the “stat” value, GSEA was performed using the GSEA software v4.3.2 (UC San Diego, USA) available at www.gsea-msigdb.org. Gene compositions of BTMs defined for humans (Li et al., 2014) were adapted to the pig genome (Bocard et al., 2021). FDR q-value below 0.05 was selected as a cutoff for significance. Normalized enrichment score (NES) and -log10(q) values were used for creating dot plots in R software with ggplot2 package. GSVA for each individual pig based on the absolute or normalized read counts was calculated using the R package GSVA (v2.0.5) with "ssgsea" method (Hänzelmann et al., 2013), allowing for a minimum size group of 15 and a maximum of 500.

### Swine leukocyte antigen (SLA) haplotyping

For SLA class I haplotyping, RNA was extracted from whole white blood cells using Trizol and NucleoSpin RNA kit (Macherey-Nagel, Germany). For SLA class II haplotyping, DNA was isolated with NucleoSpin Blood L kit (Macherey-Nagel, Germany) from 2 mL of blood collected in PAXgene Blood DNA tubes (Qiagen, Germany). DNA templates were used directly for PCR analysis. For each sample, loci-specific primer sets were applied to generate amplicons. These amplicons were barcoded individually to allow multiple samples to be analyzed on the MiSeq sequencer (Illumina, USA). For RNA templates, RT-PCR was first performed to generate cDNA. A loci-specific PCR was then performed with this cDNA, after which these products were also barcoded. Subsequently, as with DNA-templates, cDNA was analyzed on the MiSeq sequencer using the same multiplex sequencing approach. After sequencing, data from each sample was analyzed against loci-specific databases. This bioinformatics analysis enabled comparison of each sample to reference sequences to identify genetic variation per locus. BBtools suite was used for alignment.

### Statistical and correlation analyses

The analyses were performed in GraphPad Prism version 10 (GraphPad Software, USA). In order to analyze correlations between the datasets, Pearson coefficients were computed. Numbers of animals, statistical tests and p-values are specified in the corresponding figures. Dot plots were created in R software with ggplot2 package using correlation coefficient (r) and -log10(p) values. Images showing experiment and assay setups were designed in BioRender (BioRender, Canada).

## Acknowledgments

We thank Daniel Brechbühl, Katarzyna Sliz, Hans-Peter Lüthi, Jan Salchli, and Roman Troxler for excellent animal care. We thank Aurélie Godel and Markus Gerber for technical assistance. We thank Pamela Nicholson from the NGS platform of the University of Bern for library preparation and mRNA sequencing. We are grateful to Imbi Nurmoja and Olev Kalda from the Estonian Veterinary and Food Laboratory (VFL, Tartu, Estonia) for their agreement to transfer the Estonia 2014 isolate to the IVI (Mittelhäusern, Switzerland). We are grateful to Sandra Blome and Martin Beer from the Friedrich-Loeffler-Institut (FLI, Greifswald-Insel Riems, Germany) for sending the two ASFV strains (Estonia 2014 and Armenia 2008) and for the precious collaboration. We also thank Takehiro Kokuho from the Division of Transboundary Animal Disease Research at the National Institute of Animal Health (NIAH, Kodaira, Tokyo, Japan) and Ken Katsuda, Director of the National Agriculture and Food Research Organization (NARO, Ibaraki, Japan) for providing the IPKM cell line. Finally, we would like to thank the members of the ASF-RASH consortium for fruitful discussions and continued support. This research was partially supported by the Veterinary Food Safety and Veterinary Office (Grant No. 1.21.12) under the umbrella of the ICRAD ERANET ASF-RASH, co-funded European Union’s Horizon 2020 research and innovation program (Grant Agreement No. 862605). Additional support was provided by the Staatssekretariat für Bildung, Forschung und Innovation (SBFI; Project No. 23.00633), under the Horizon 2020 research and innovation program ASFaVIP (Grant Agreement No. 101136676).

