## Supplementary figures for "Correlates of protection against African swine fever virus identified by a systems immunology approach"

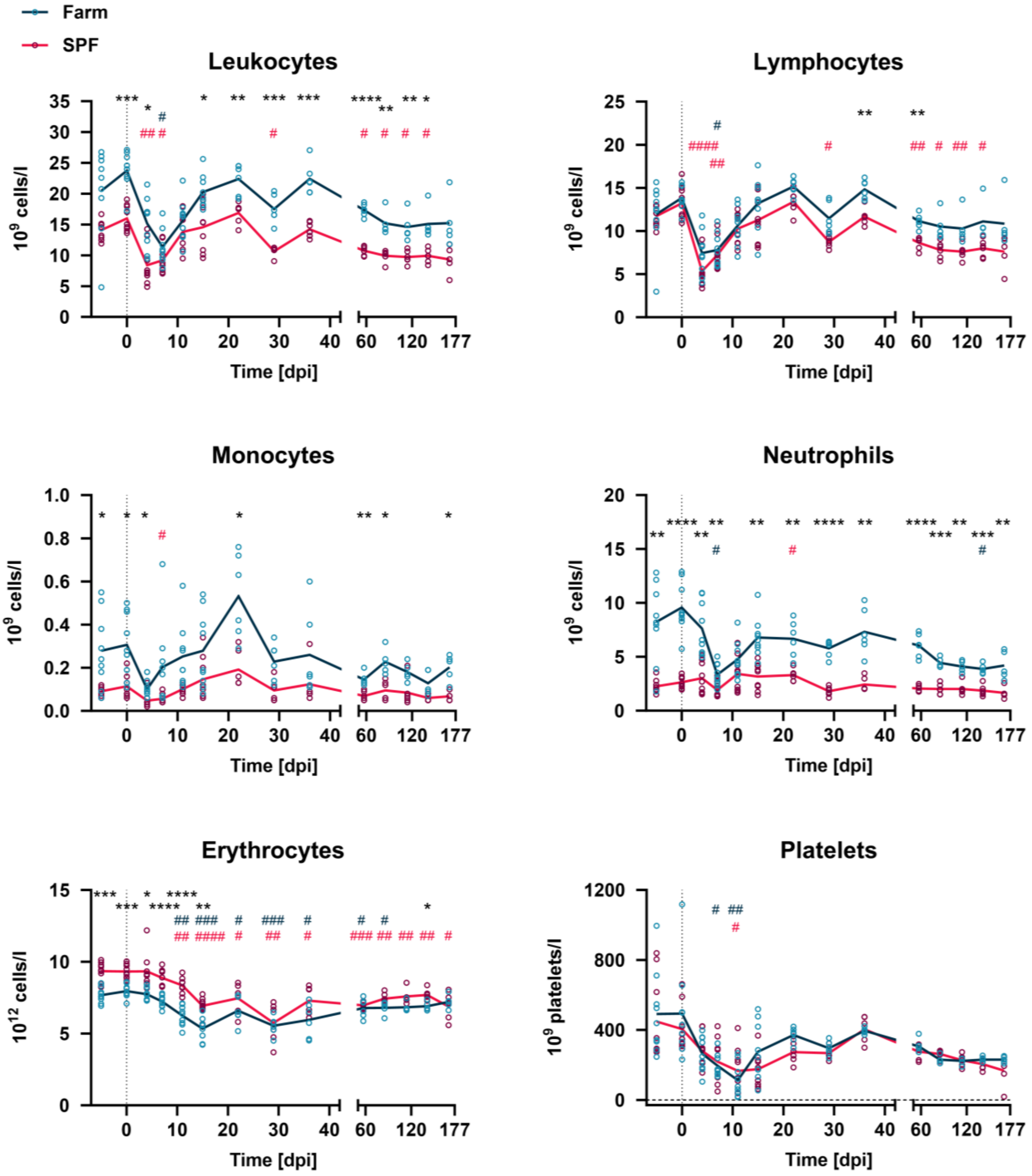


**Supplementary Fig. 1. Hematologic profiles after immunization with the Estonia 2014 strain.** Leukocyte, erythrocyte, and platelet counts with data points representing individual animals and lines showing the means of farm and SPF groups. There were n = 9 pigs in each group. Differences to baseline measurements (day -5) for farm and SPF groups were analyzed by mixed-effects analysis with Dunnett's multiple comparisons test (blue hashtags - farm group, pink hashtags - SPF group); $^{\#}p < 0.05$; $^{\#\#}p < 0.01$; $^{\#\#\#}p < 0.001$; $^{\#\#\#\#}p < 0.0001$. Differences between farm and SPF groups were analyzed at each time point by unpaired t-test with Holm-Šidák's correction for multiple comparisons; $^{*}p < 0.05$; $^{**}p < 0.01$; $^{***}p < 0.001$; $^{****}p < 0.0001$.

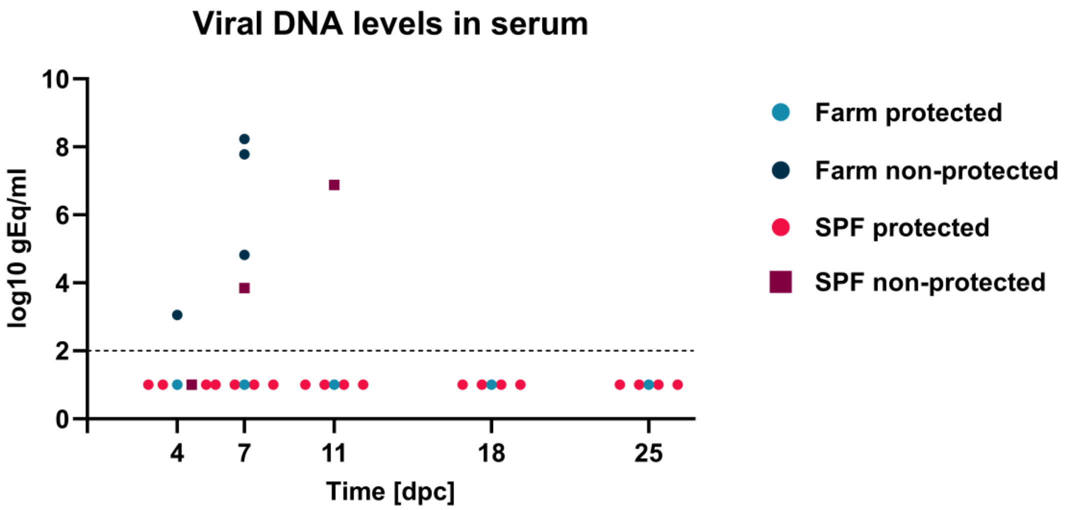


**Supplementary Fig. 2. Viral DNA levels in serum after challenge.** Quantification was performed by qPCR. Data points represent values for individual animals remaining in the experiment at various time points post-challenge. At 4 and 7 dpc, n = 5 pigs in farm and SPF groups, at 11 dpc, n = 2 in farm group and n = 5 in SPF group, and at 18 and 25 dpc, n = 2 in farm group and n = 4 in SPF group.

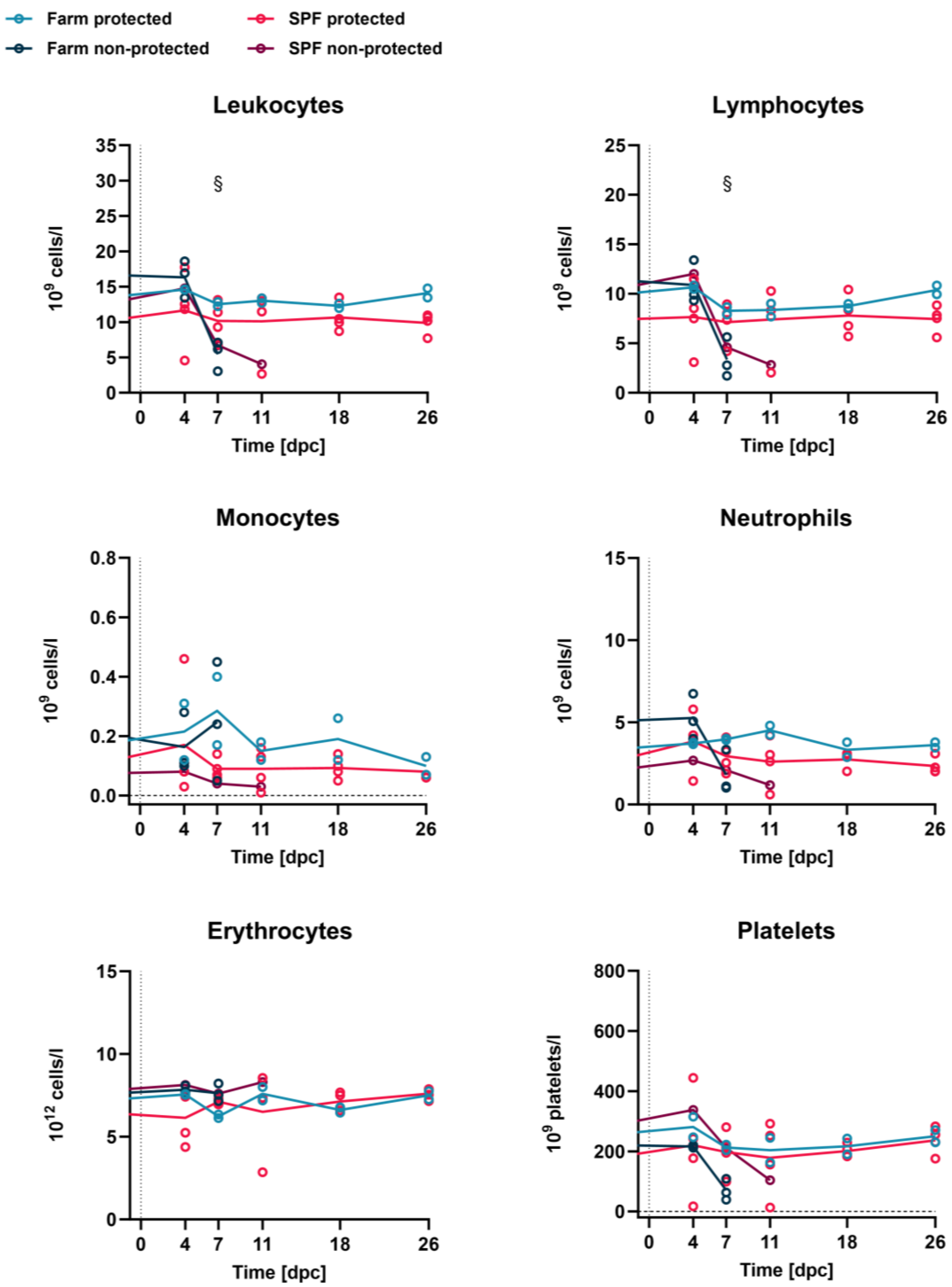


**Supplementary Fig. 3. Hematologic profiles after challenge with the Armenia 2008 strain.** Leukocyte, erythrocyte, and platelet counts of farm and SPF pigs. Data points represent values for individual animals, lines show the means of the groups. At 4 and 7 dpc, n = 5 pigs in farm and SPF groups, at 11 dpc, n = 2 in farm group and n = 5 in SPF group, and at 18 and 26 dpc, n = 2 in farm group and n = 4 in SPF group. Differences between protected and non-protected animals were analyzed by unpaired t-test with Holm-Šídák's correction for multiple comparisons; $^{§}p < 0.05$.

**A**

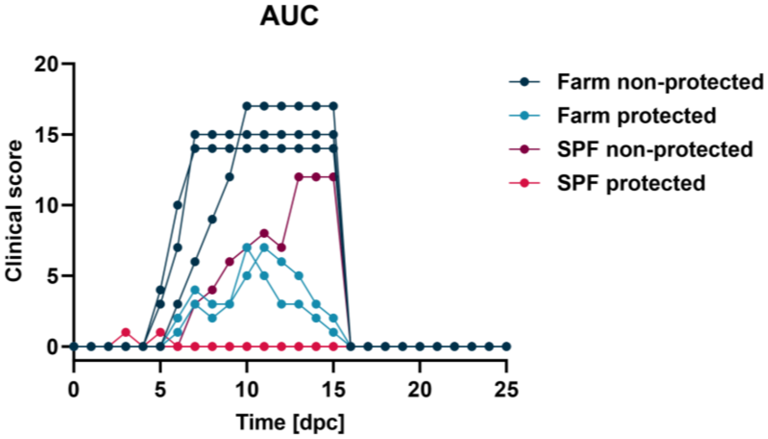


**B**

| Pig | AUC | Pig | AUC |
|---|---|---|---|
| #13 | 132 | #1963 | 0 |
| #14 | 40 | #1966 | 0 |
| #15 | 140 | #1967 | 71 |
| #16 | 30 | #1973 | 1 |
| #17 | 145 | #1974 | 1 |

**Supplementary Fig. 4. Clinical scores values for AUC calculations.** (A) The clinical scores post-challenge, used to calculate the area under the curve (AUC) for each animal, are shown. The lines were prolonged until 16 dpc for the animals reaching clinical end points to obtain an expected minimum AUC. (B) AUC values for farm and SPF pigs used for correlations.

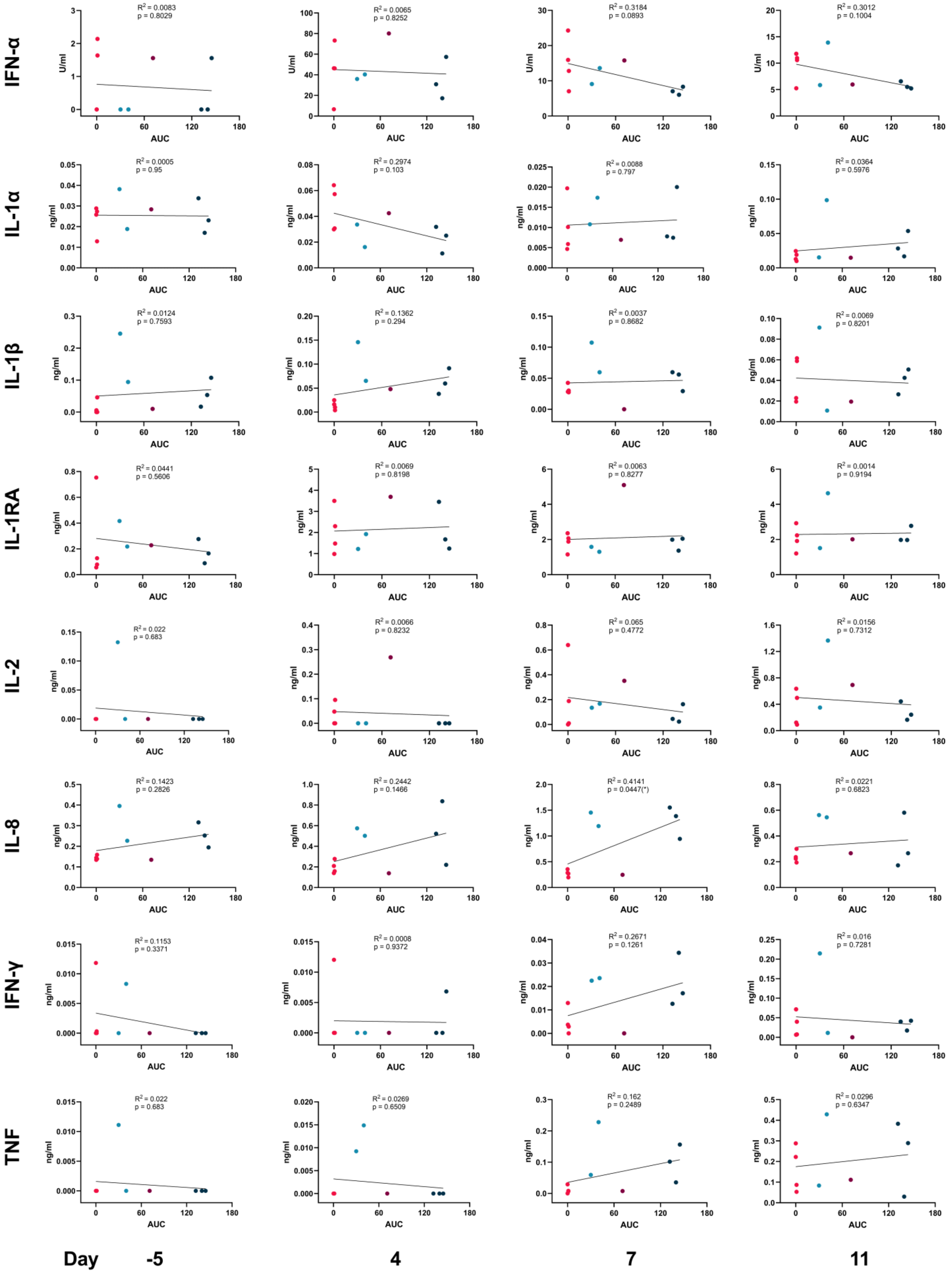


**Supplementary Fig.5. Correlation of serum cytokine levels post-immunization with clinical outcomes (AUCs).** For each group, n = 5 pigs were included in the analysis.

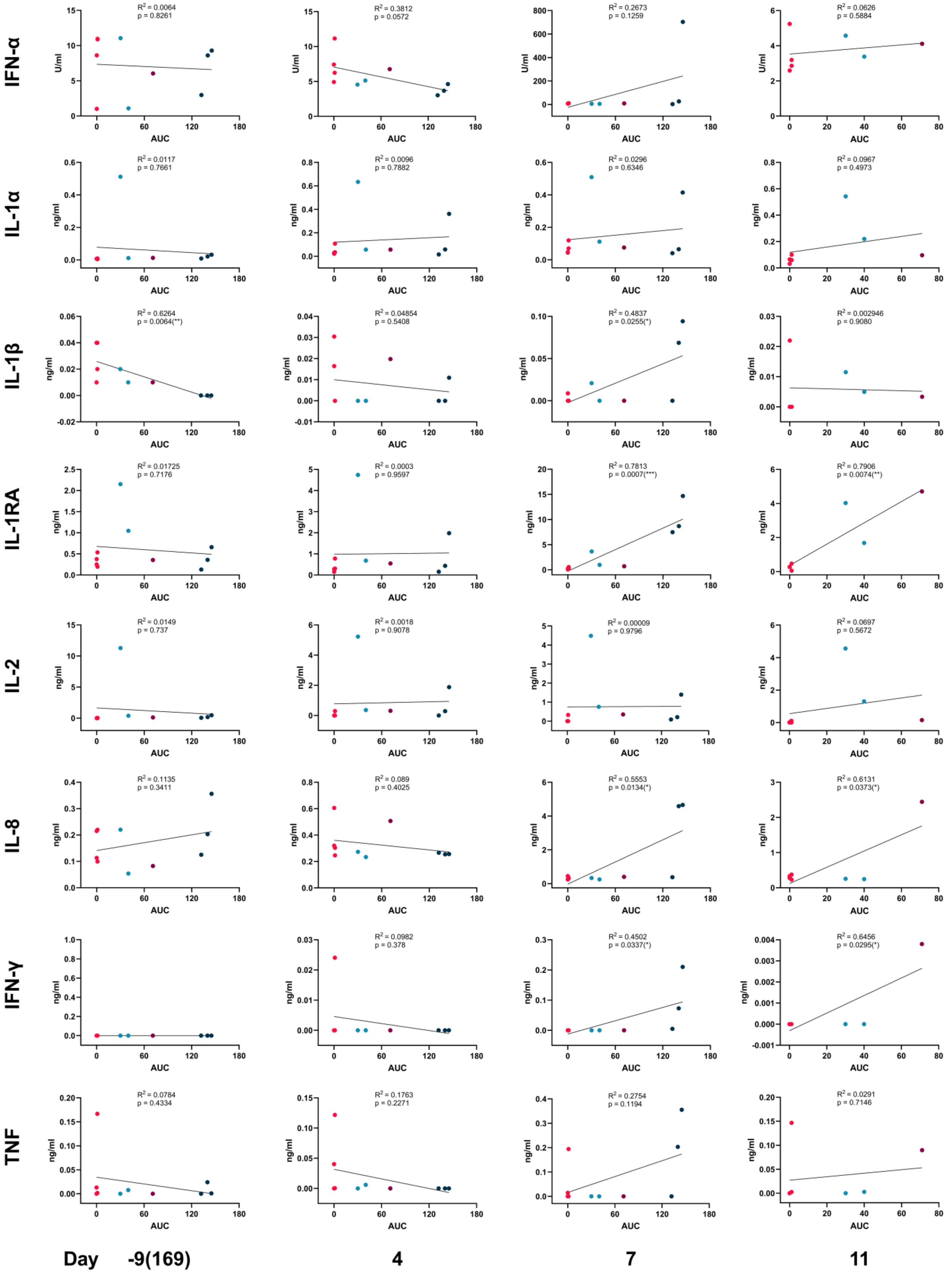


**Supplementary Fig. 6. Correlation of serum cytokine levels post-challenge with clinical outcomes (AUCs).** At -9, 4 and 7 dpc, n = 5 pigs in farm and SPF groups, and at 11 dpc, n = 2 in farm group and n = 5 in SPF group.

| Antigen | Clone | Species/Isotype | Fluorochrome | Source | Cat No. |
|---|---|---|---|---|---|
| CD3 | PPT3 | Mouse IgG1 | - | Hybridomas | - |
| CD4 | 74-12-4 | Mouse IgG2b | - | Hybridomas | - |
| CD8α | 76-2-11 | Mouse IgG2a | PE | BD Pharmingen | 559584 |
| CD8β | PG164A | Mouse IgG2a | - | WSU | PG164A |
| δ-TCR* | PGBL22A | Mouse IgG1 | - | WSU | PGBL22A |
| IFN-γ | P2G10 | Mouse IgG1 | PerCP-Cy5.5 | BD Pharmingen | 561481 |
| TNF-α | MAb11 | Mouse IgG1 | AF647 | BioLegend | 502916 |
| IgG1 | Polyclonal | Goat IgG | APC/CY7 | SouthernBiotech | 1070-19 |
| IgG2b | Polyclonal | Goat IgG | AF488 | Invitrogen | A-21141 |
| IgG2a | Polyclonal | Goat IgG | PE/Cy7 | Abcam | ab130787 |

**Supplementary Table 1. List of antibodies used for flow cytometry.** *The antibody against δ-TCR chain was coupled to biotin using Zenon Mouse IgG1 labeling kit (Z25052, Invitrogen, USA). Streptavidin coupled with BV421 was used as a conjugate (563259, BD Horizon, USA).

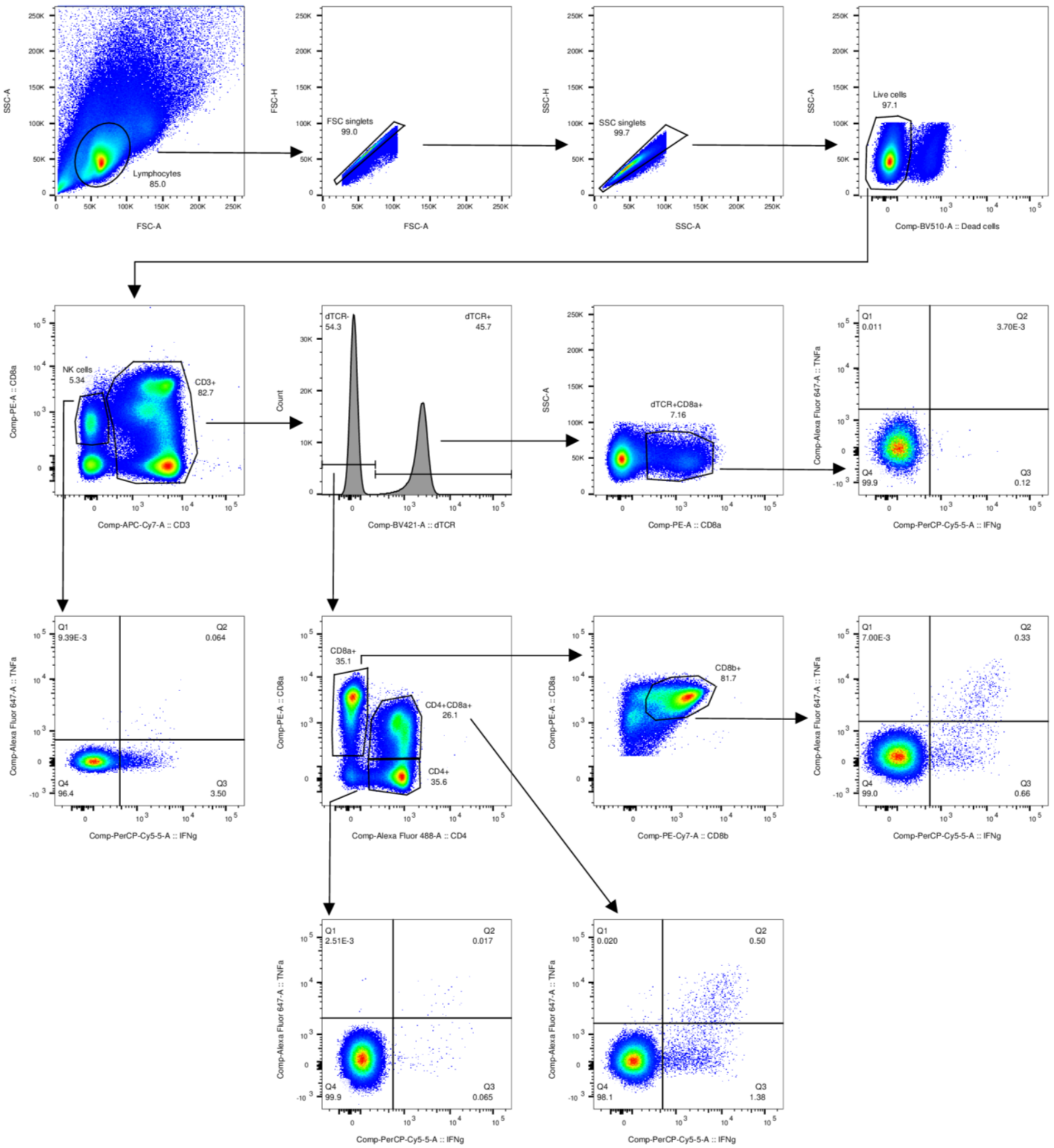


**Supplementary Fig. 7. Gating strategy for analysis of intracellular cytokine responses following *in vitro* restimulation of freshly collected PBMCs with live ASFV.**

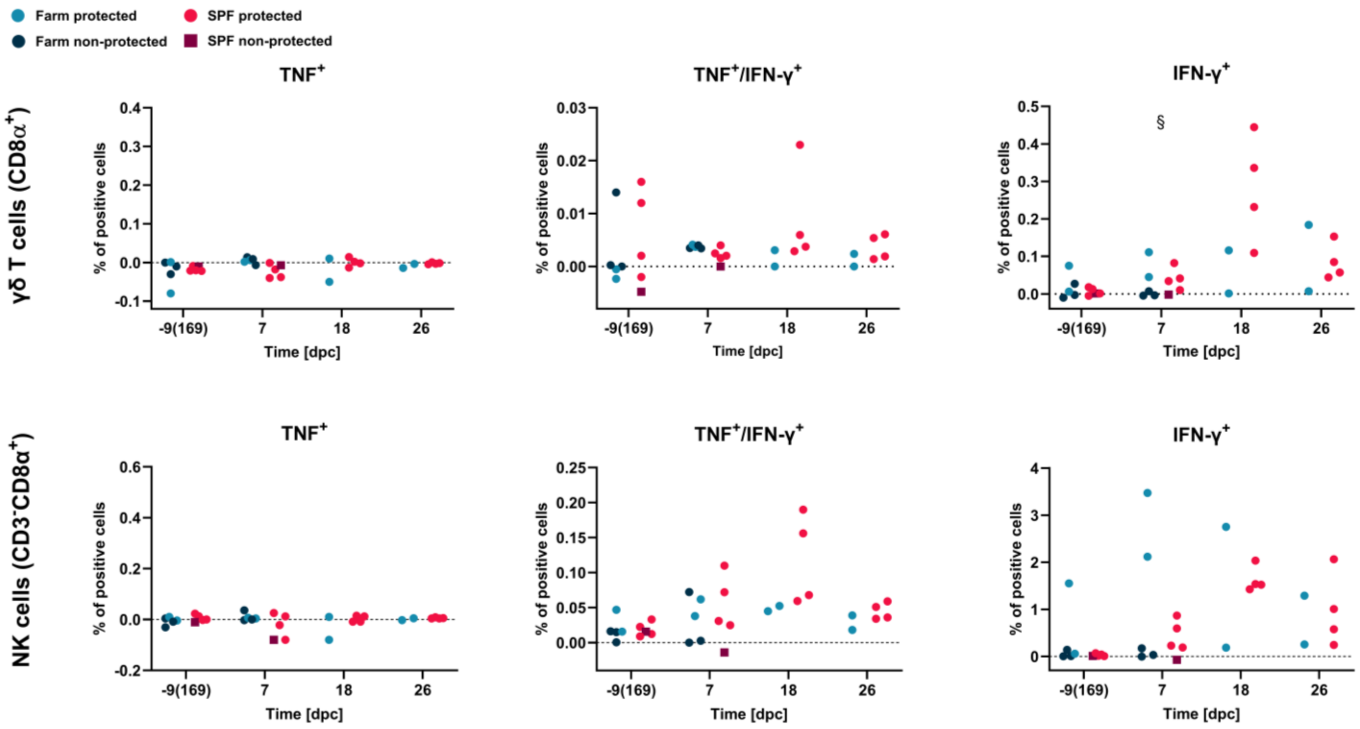


**Supplementary Fig. 8. Cellular immune responses from effector γδ T cells and NK cells during the challenge phase.** Freshly collected PBMCs were restimulated with ASFV. Supernatant from mock-infected macrophages served as negative control, obtained percentages were subtracted. Data points represent values for individual animals. At -9 and 7 dpc, n = 5 pigs in farm and SPF groups, and at 18 and 26 dpc, n = 2 in farm group and n = 4 in SPF group. Differences between protected and non-protected animals were analyzed by unpaired t-test with Holm-Šídák's correction for multiple comparisons; $^{§}p < 0.05$.

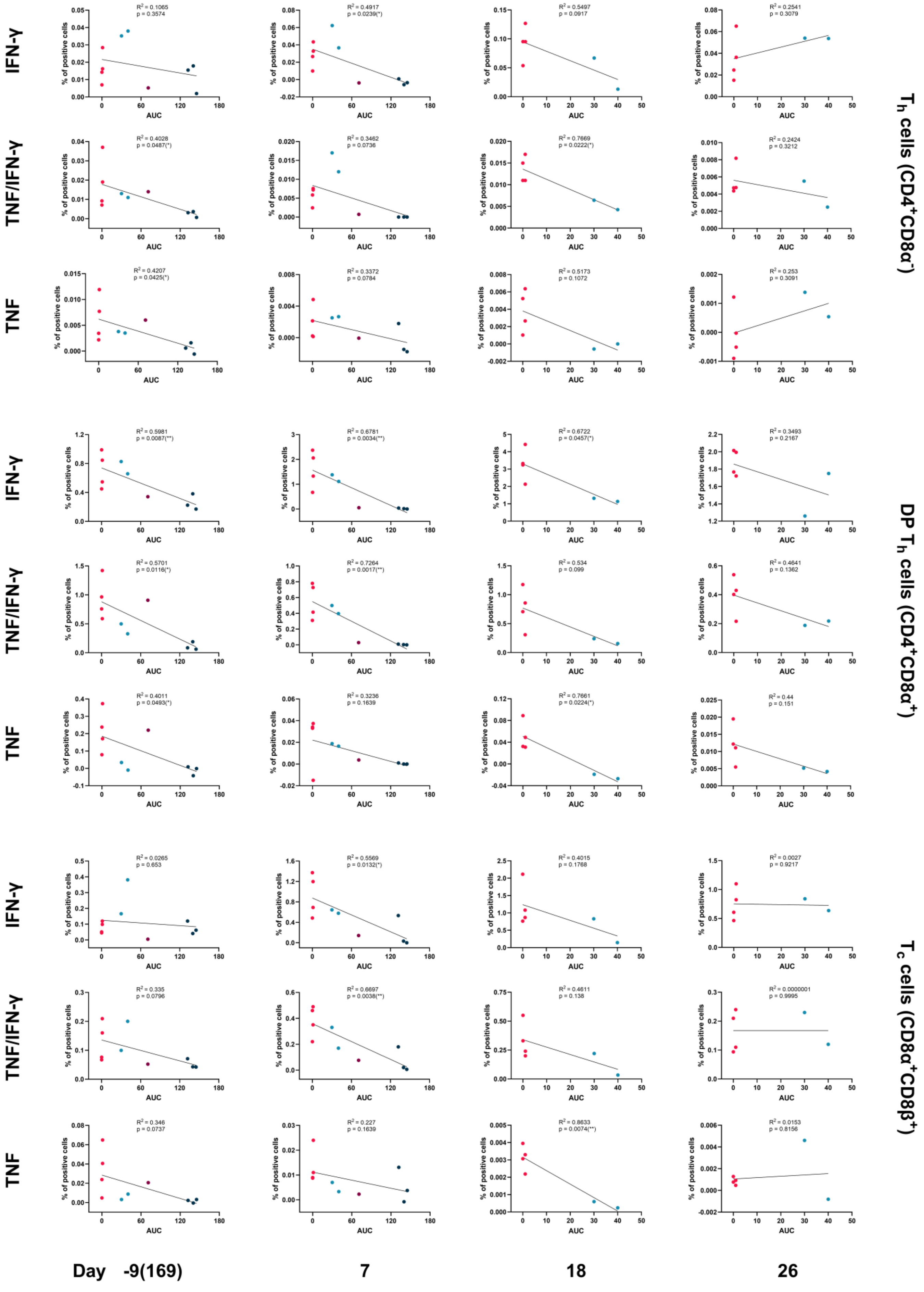


**Supplementary Fig. 9. Correlation of cytokine responses in three cell subsets post-challenge with clinical outcomes (AUCs).** At -9 and 7 dpc, n = 5 pigs in farm and SPF groups, and at 18 and 26 dpc, n = 2 in farm group and n = 4 in SPF group.

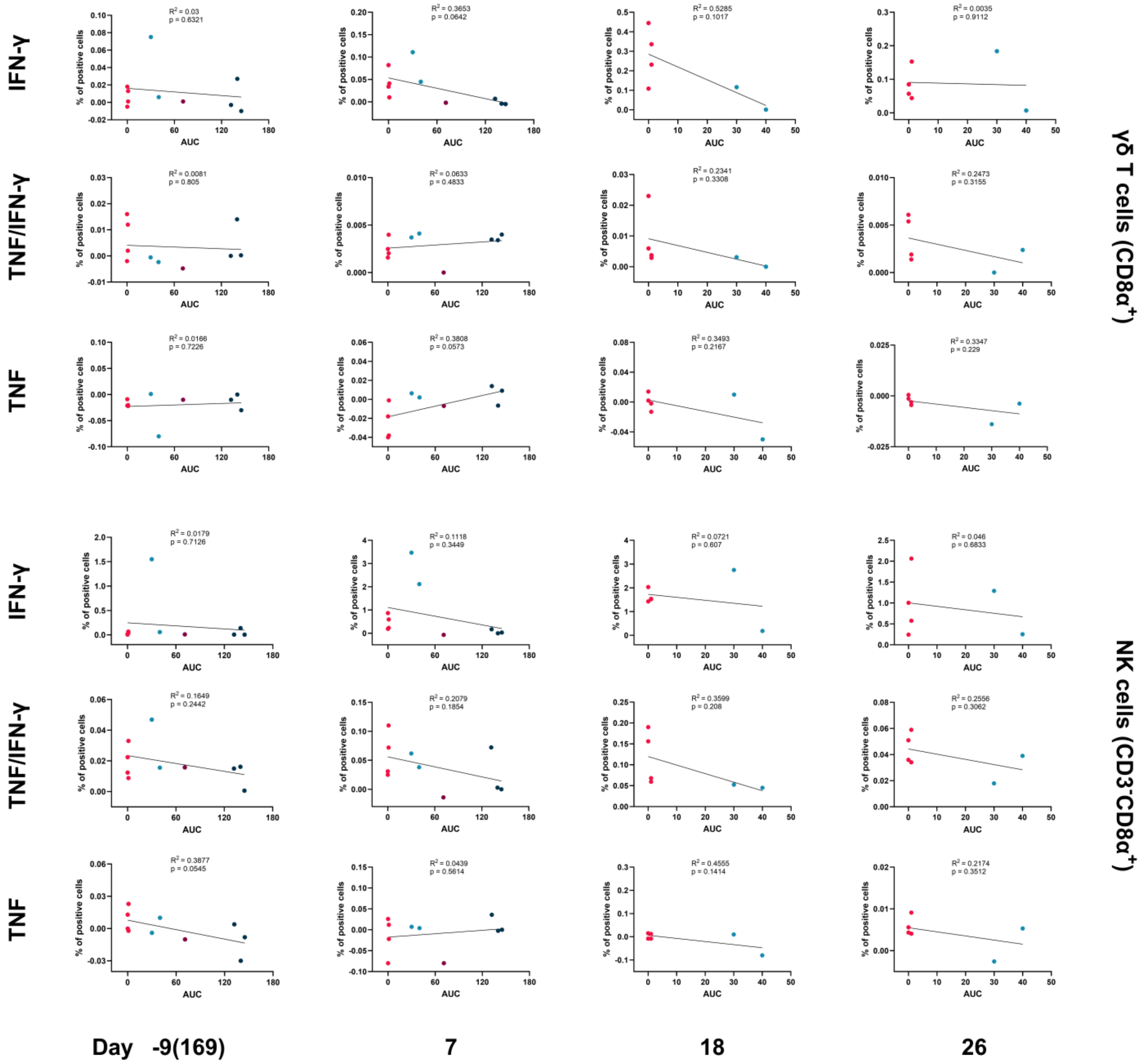


**Supplementary Fig. 10. Correlation of cytokine responses in two cell subsets post-challenge with clinical outcomes (AUCs).** At -9 and 7 dpc, n = 5 pigs in farm and SPF groups, and at 18 and 26 dpc, n = 2 in farm group and n = 4 in SPF group.

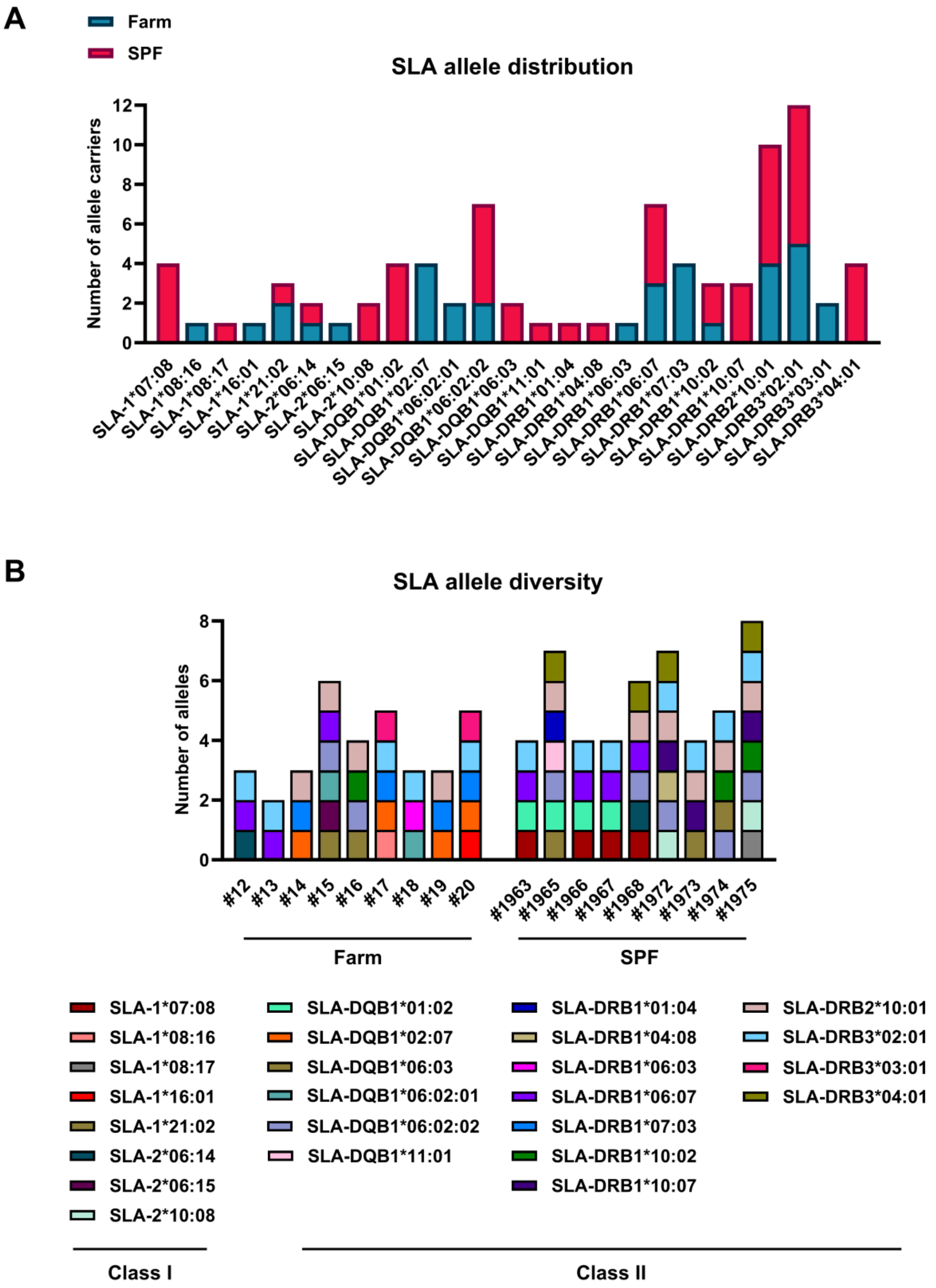


**Supplementary Fig. 11. Swine leukocyte antigen (SLA) haplotyping.** (A) Distribution of alleles for MHC (swine leukocyte antigen, SLA) class I and class II in farm and SPF pigs. (B) Allele diversity and composition of haplotypes in farm and SPF pigs.

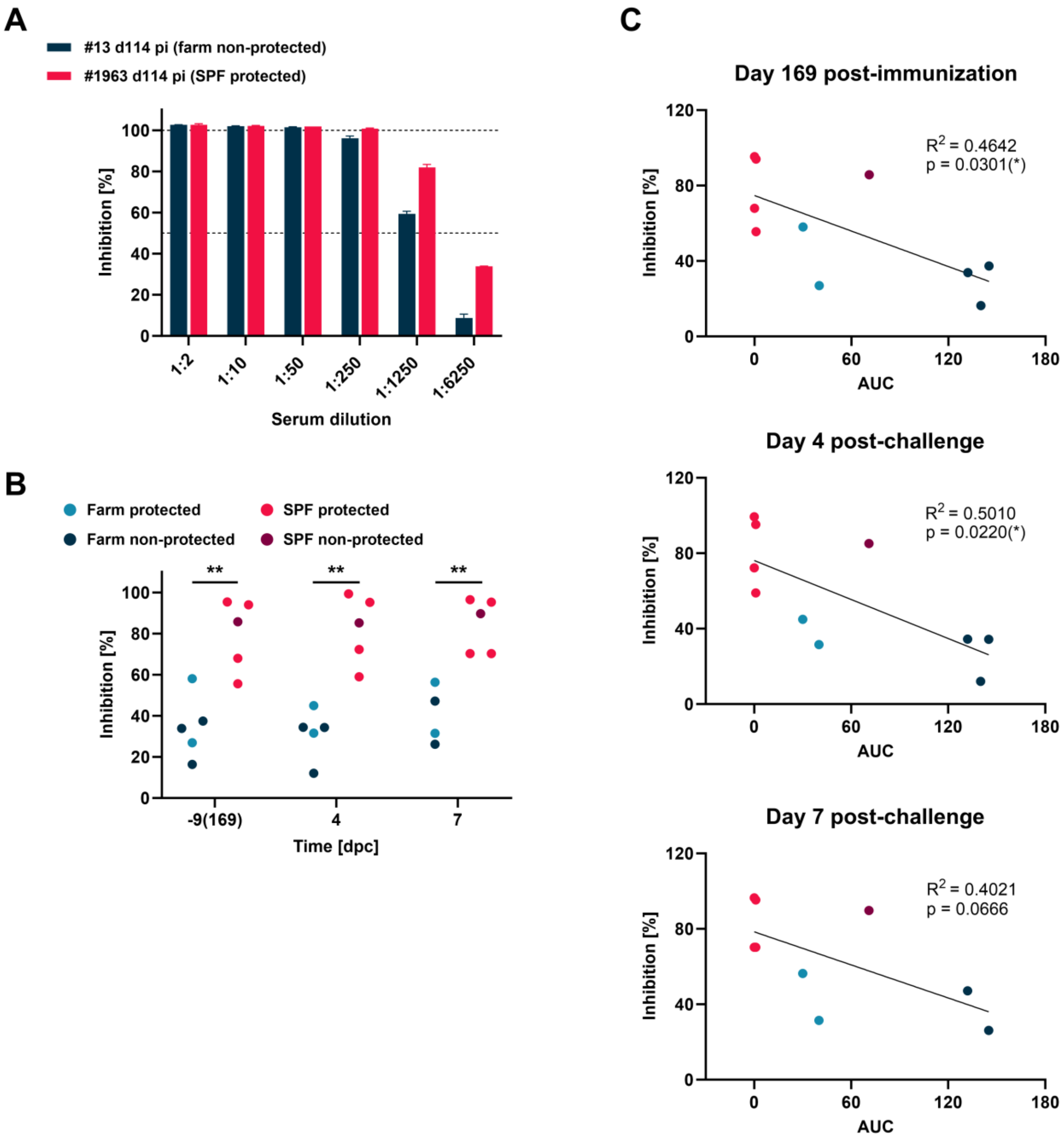


**Supplementary Fig. 12. Correlation analysis between anti-p72 antibody titers and protection.** (A) Two serum samples from a randomly selected protected SPF and a non-protected farm pig were serially diluted to define by competitive ELISA the non-saturating concentration estimated to be between 20% and 80% inhibition. The dilution factor of 1:1250 was chosen to test all serum samples. (B) Semiquantitative antibody levels expressed as percent inhibition in samples collected before and after challenge. (C) Correlation of antibody levels expressed as % inhibition with clinical outcomes. Both farm and SPF groups had n = 5 pigs each. (B) Differences between farm and SPF groups were analyzed at each time point by unpaired t-test with Holm-Šidák's correction for multiple comparisons; **p < 0.01.

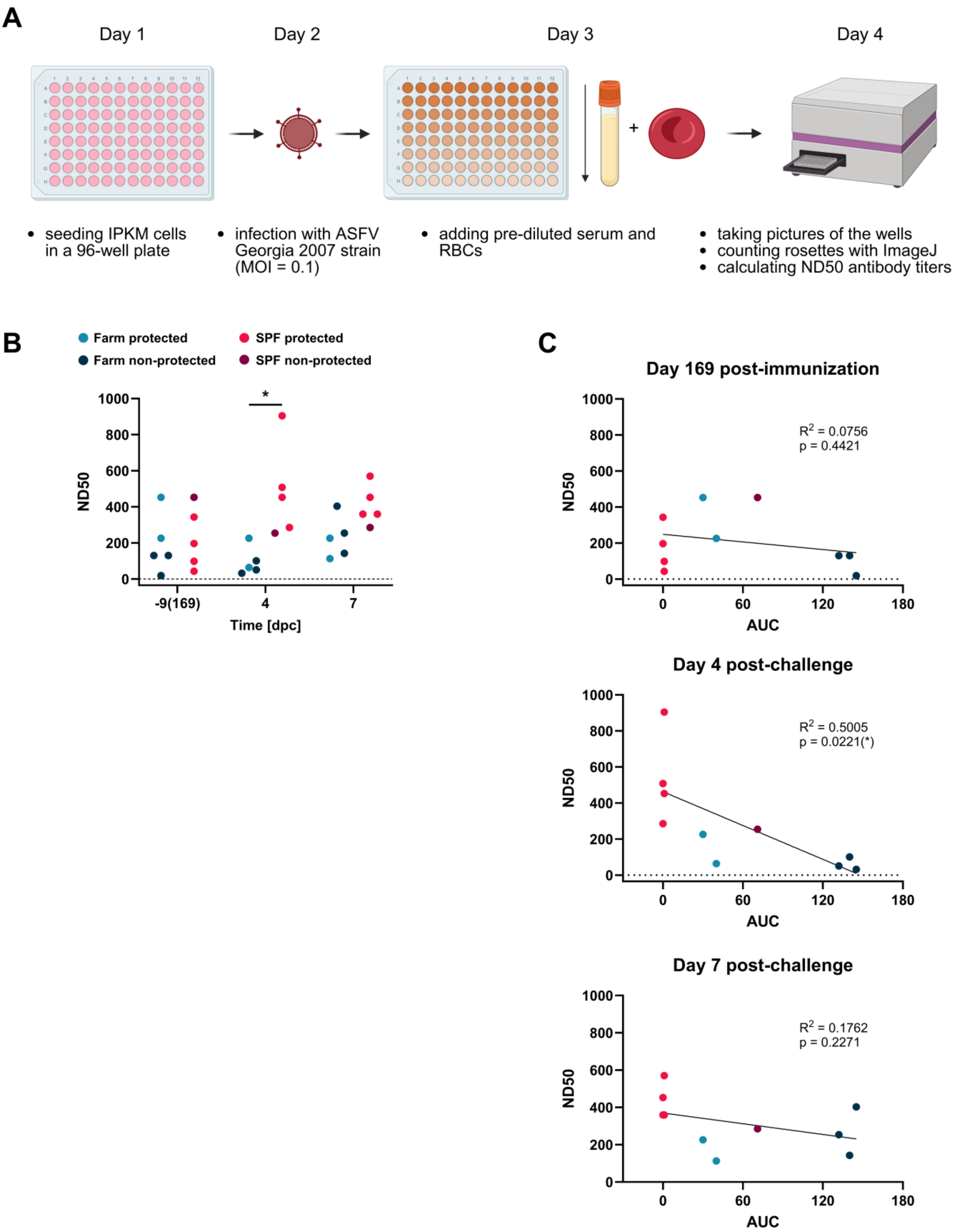


**Supplementary Fig. 13. Correlation analysis between hemadsorption-inhibiting antibody titers and protection.** (A) Workflow of hemadsorption inhibition assay (HADIA) for determining antibody titers. (B) Comparison of antibody titers in samples collected before and after challenge. (C) Correlation of antibody titers with clinical outcomes. Both farm and SPF groups had n = 5 pigs each. (B) Differences between farm and SPF groups were analyzed at each time point by unpaired t-test with Holm-Šidák's correction for multiple comparisons; *p < 0.05.

A

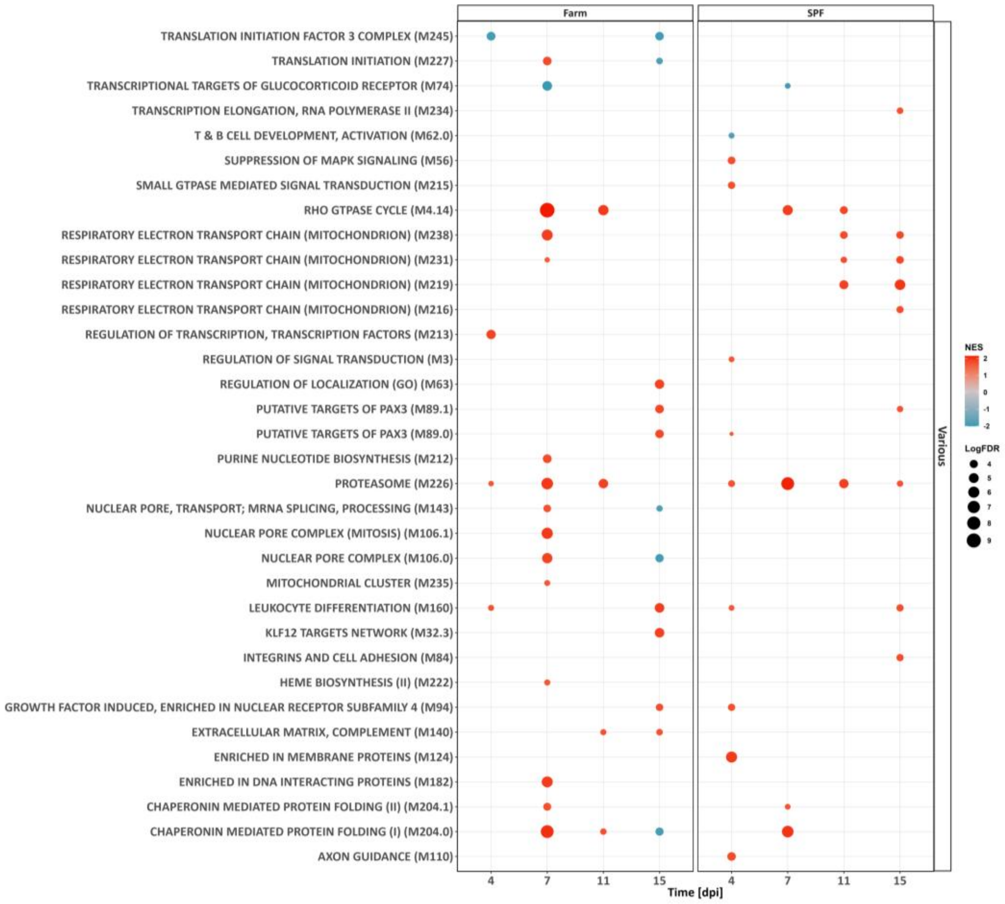


B

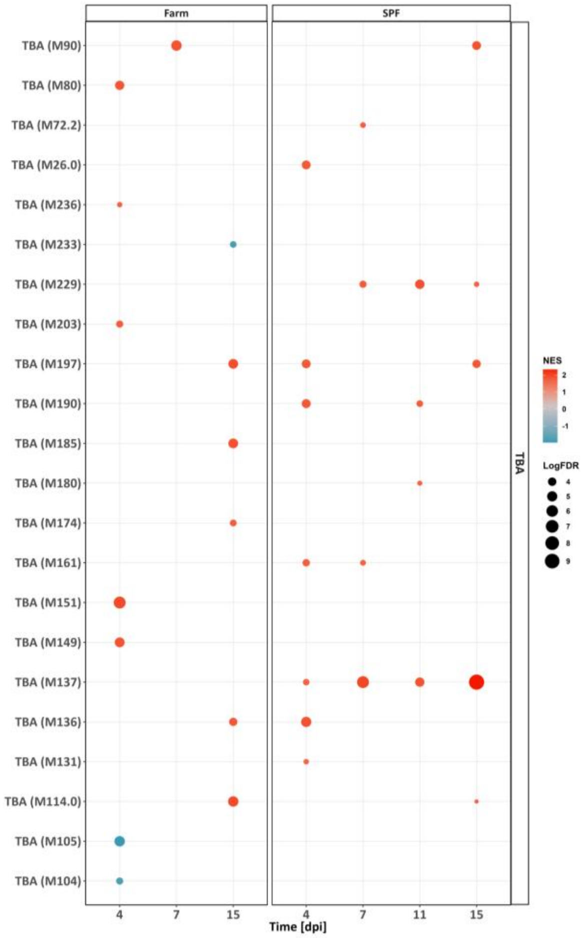


**Supplementary Fig. 14. Blood leukocyte transcriptomic profiles after immunization with the Estonia 2014 strain.** For each group of animals, comparisons were made against the baseline measurements (day 0). Pre-ranked list of DEGs was subjected to GSEA using BTMs as gene sets. (A, B) Dot plots show non-classified and to-be-annotated (TBA) BTMs, respectively. Dot size depicts the q-value (FDR), while color represents the normalized enrichment score (NES). FDR < 0 .05 was selected as a cutoff.

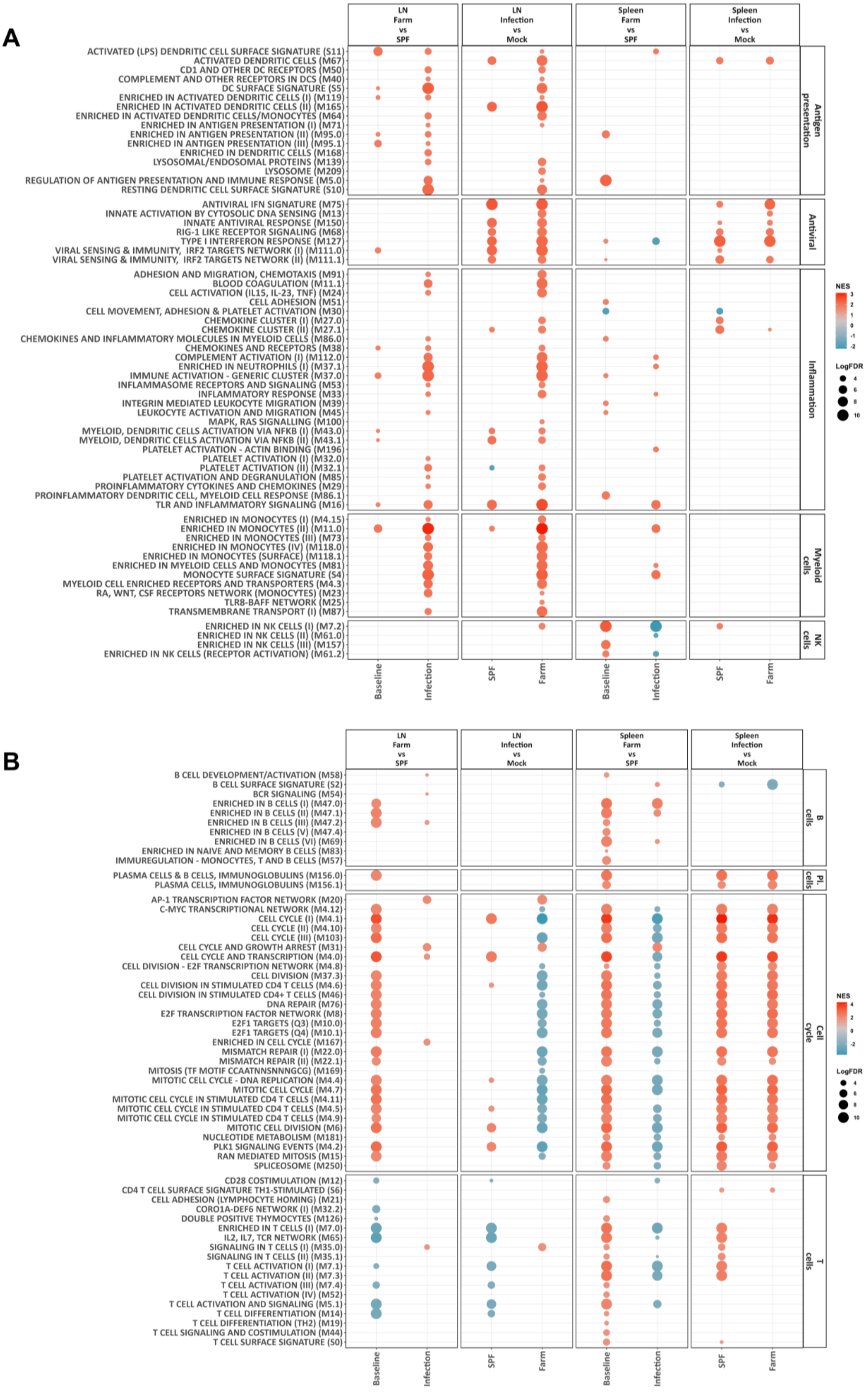


**Supplementary Fig. 15. Lymphoid tissue transcriptomic profiles after immunization with the Estonia 2014 strain.** Gastrohepatic lymph nodes (LNs) and spleens were collected at 17 dpi. For each group of animals, comparisons were made against the mock group. In addition, farm and SPF pigs were compared between each other. Pre-ranked list of DEGs was subjected to GSEA using BTMs as gene sets. (A, B) Dot plots show innate and adaptive BTMs, respectively. Dot size depicts the q-value (FDR), while color represents the normalized enrichment score (NES). FDR < 0.05 was selected as a cutoff.

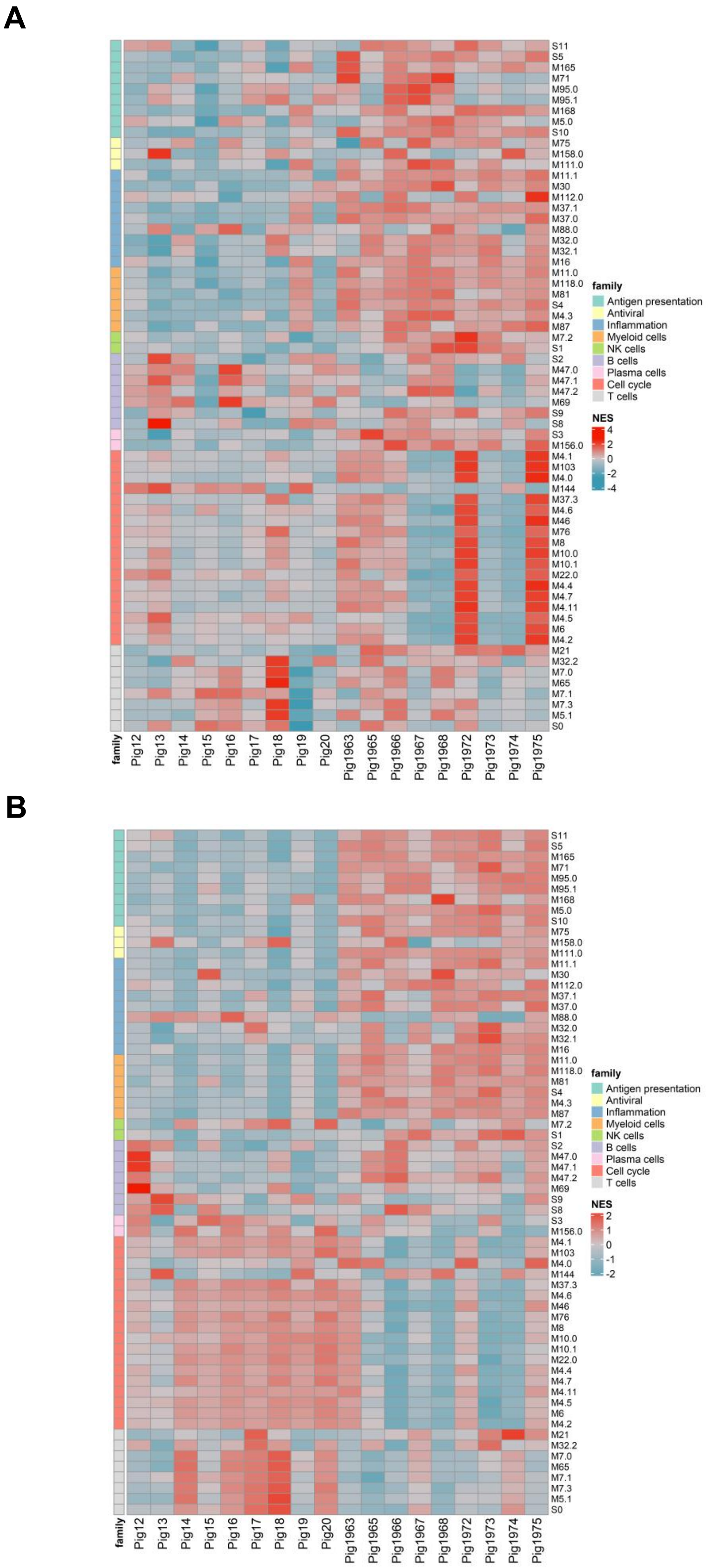


**Supplementary Fig. 16. Heatmaps showing relative BTM expression in blood leukocytes at 4 dpi (A) and 7 dpi (B).** Gene expression counts were obtained after bulk RNA-seq and normalized using day 0 values. GSVA was then performed with BTM gene sets to calculate NES values displayed on the heatmaps. BTM families are indicated by the color codes (left side) and BTM identifiers are listed on the right side. Pigs 12–20: farm group; pigs 1963–1975: SPF group.

A

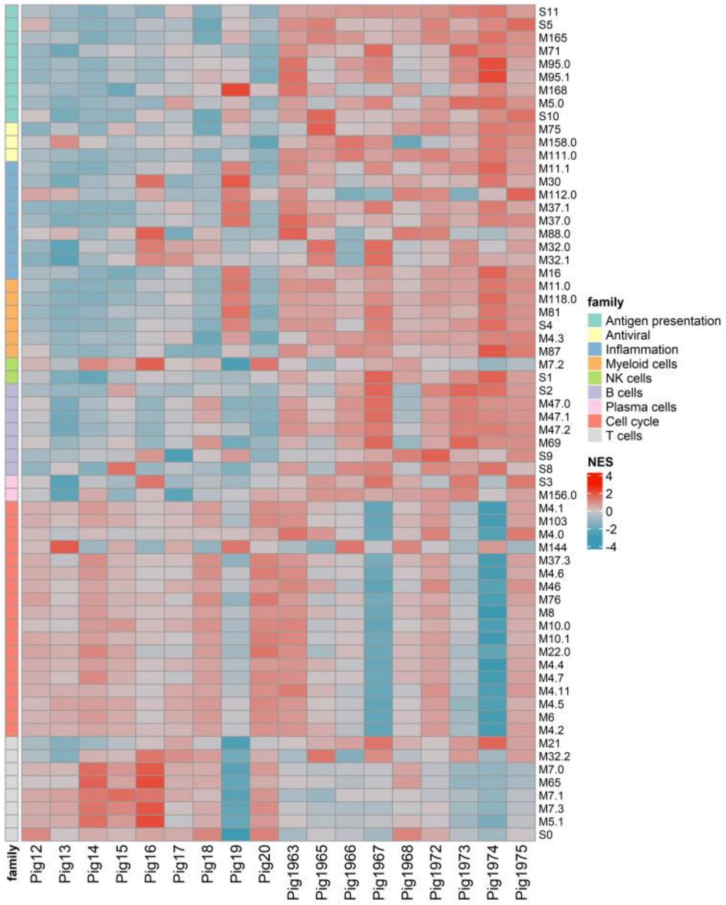


B

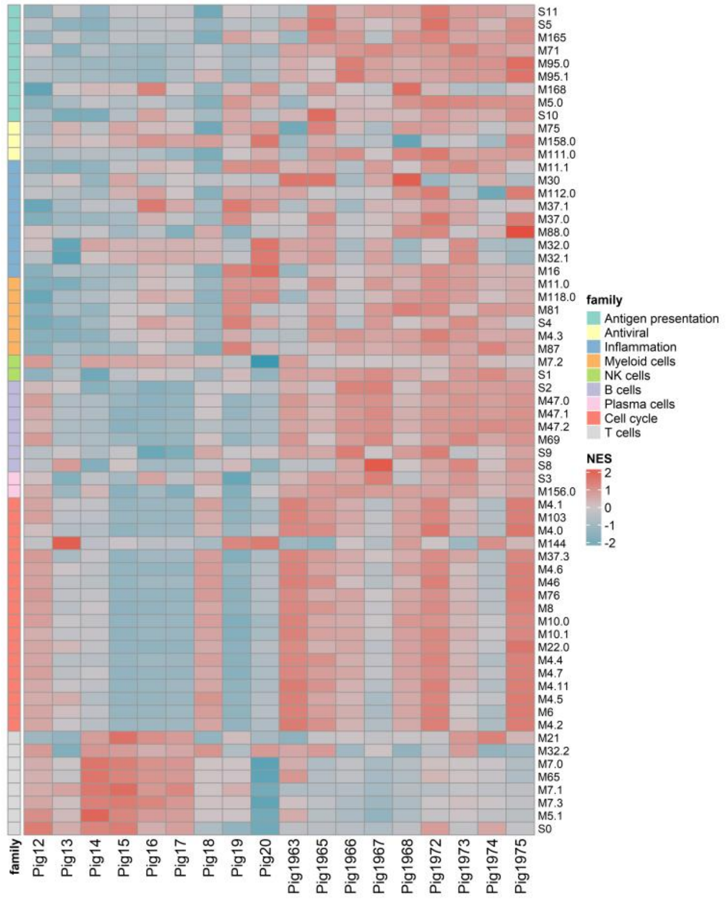


**Supplementary Fig. 17. Heatmaps showing relative BTM expression in blood leukocytes at 11 dpi (A) and 15 dpi (B).** Gene expression counts were obtained after bulk RNA-seq and normalized using day 0 values. GSVA was then performed with BTM gene sets to calculate NES values displayed on the heatmaps. BTM families are indicated by the color codes (left side) and BTM identifiers are listed on the right side. Pigs 12–20: farm group; pigs 1963–1975: SPF group.

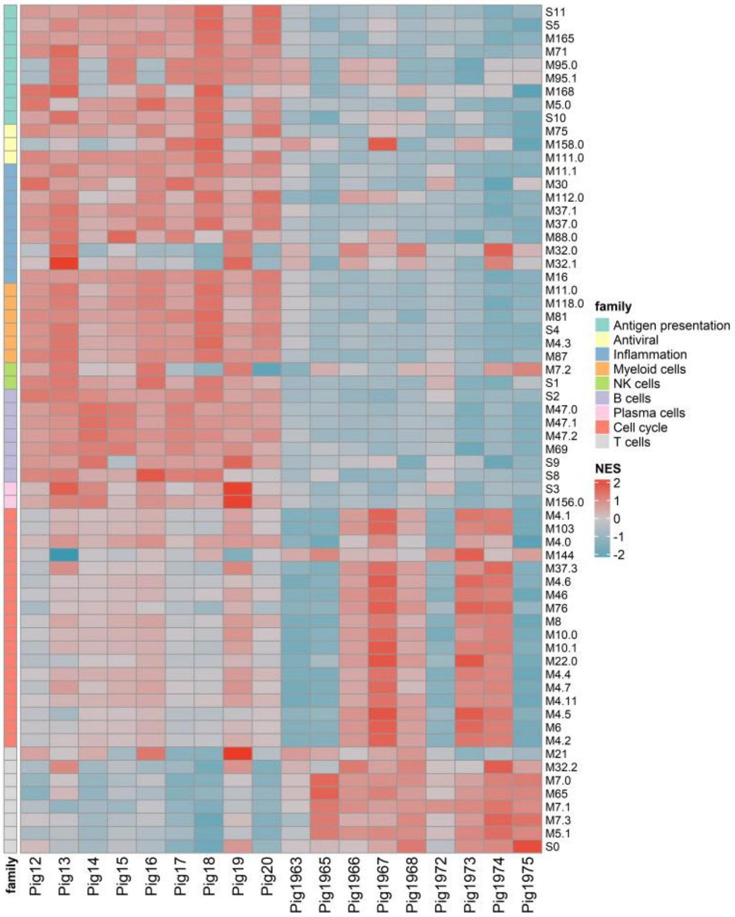


**Supplementary Fig. 18. Heatmap showing absolute BTM expression in blood leukocytes at baseline (day 0).** Gene expression counts were obtained after bulk RNA sequencing and used without normalization. GSVA was then performed with BTM gene sets to calculate NES values displayed on the heatmap. BTM families are indicated by the color codes (left side) and BTM identifiers are listed on the right side. Pigs 12–20: farm group; pigs 1963–1975: SPF group.

A

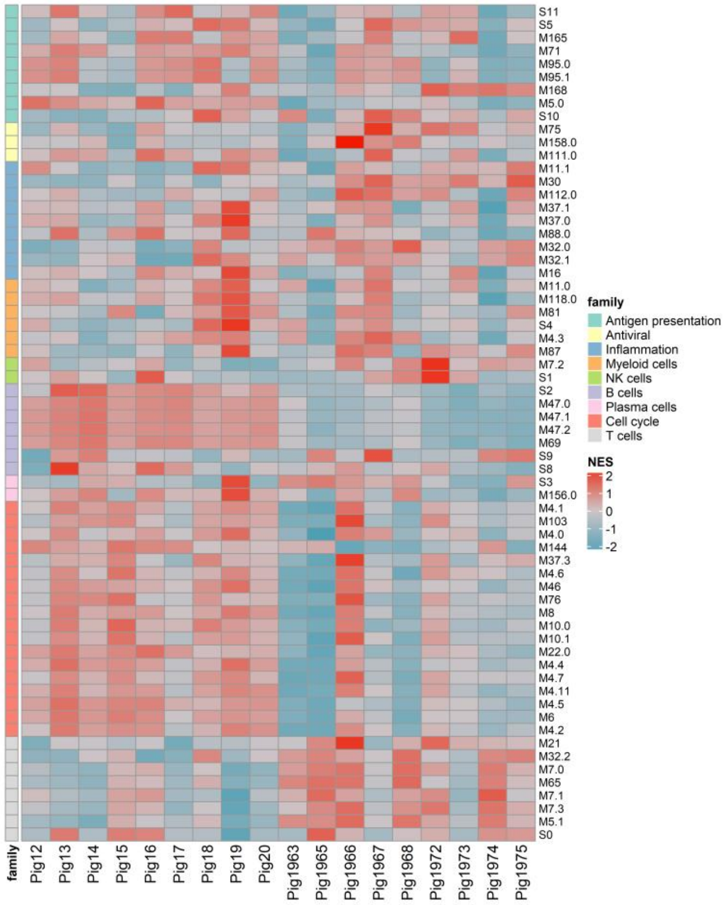


B

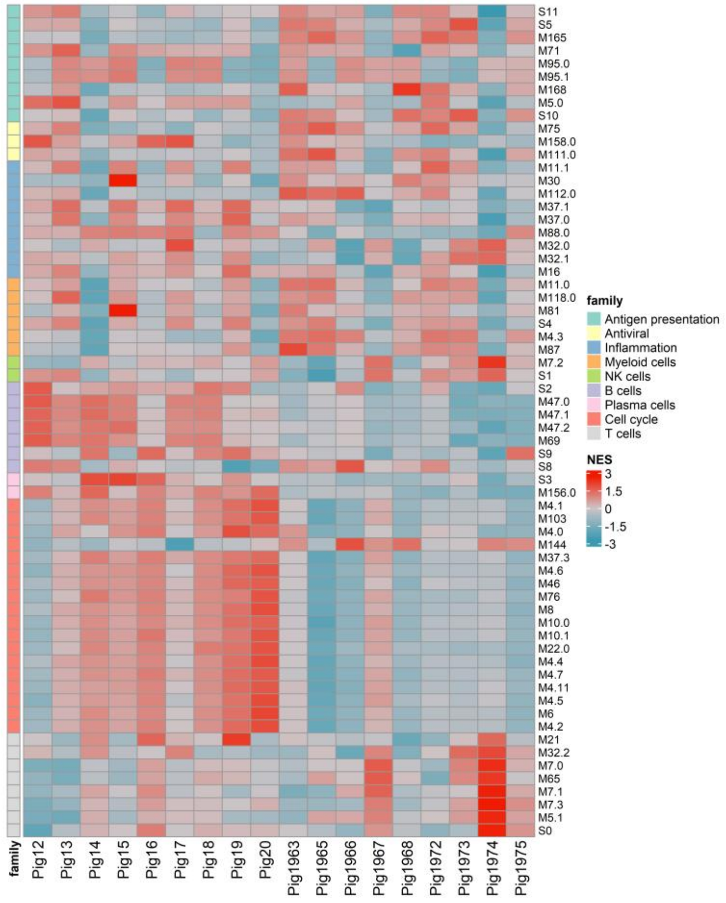


**Supplementary Fig. 19. Heatmaps showing absolute BTM expression in blood leukocytes at 4 dpi (A) and 7 dpi (B).** Gene expression counts were obtained after bulk RNA-seq and and used without normalization to day 0 values. GSVA was then performed with BTM gene sets to calculate NES values displayed on the heatmaps. BTM families are indicated by the color codes (left side) and BTM identifiers are listed on the right side. Pigs 12–20: farm group; pigs 1963–1975: SPF group.

A

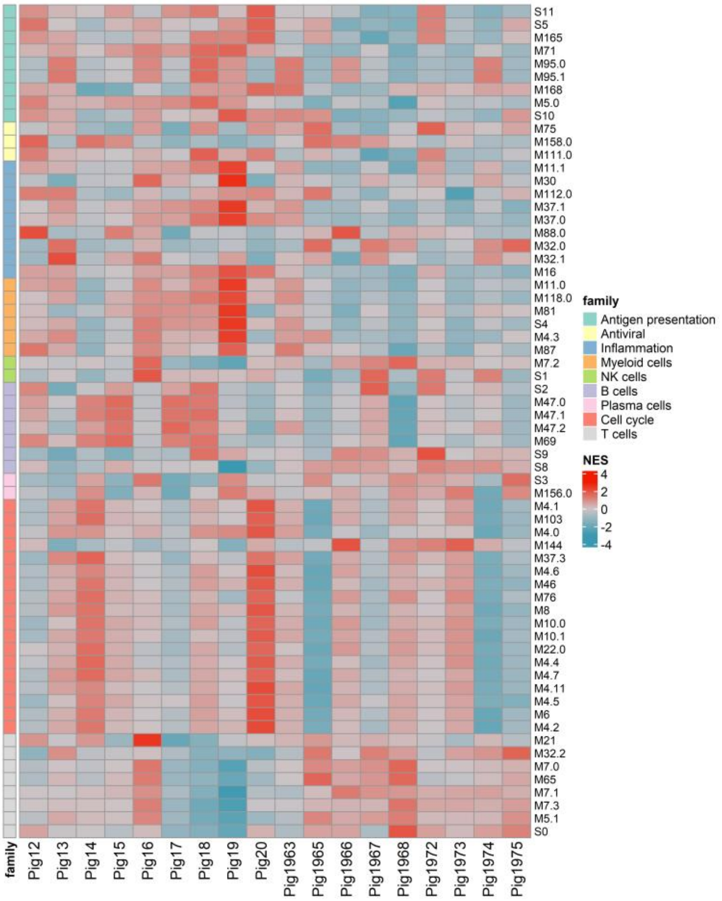


B

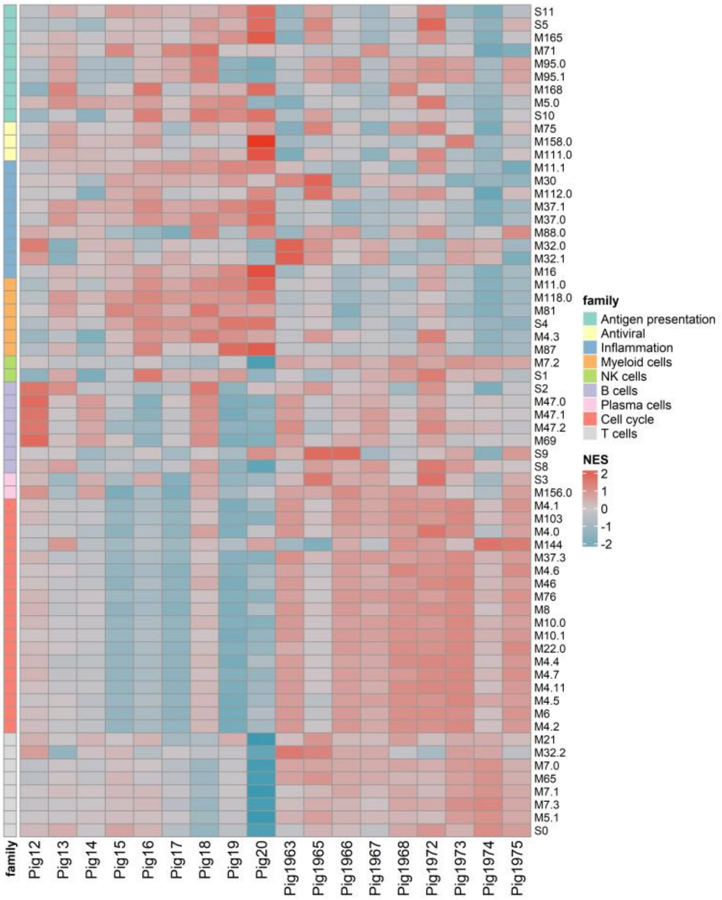


**Supplementary Fig. 20. Heatmaps showing absolute BTM expression in blood leukocytes at 11 dpi (A) and 15 dpi (B).** Gene expression counts were obtained after bulk RNA-seq and and used without normalization to day 0 values. GSVA was then performed with BTM gene sets to calculate NES values displayed on the heatmaps. BTM families are indicated by the color codes (left side) and BTM identifiers are listed on the right side. Pigs 12–20: farm group; pigs 1963–1975: SPF group.

A

B

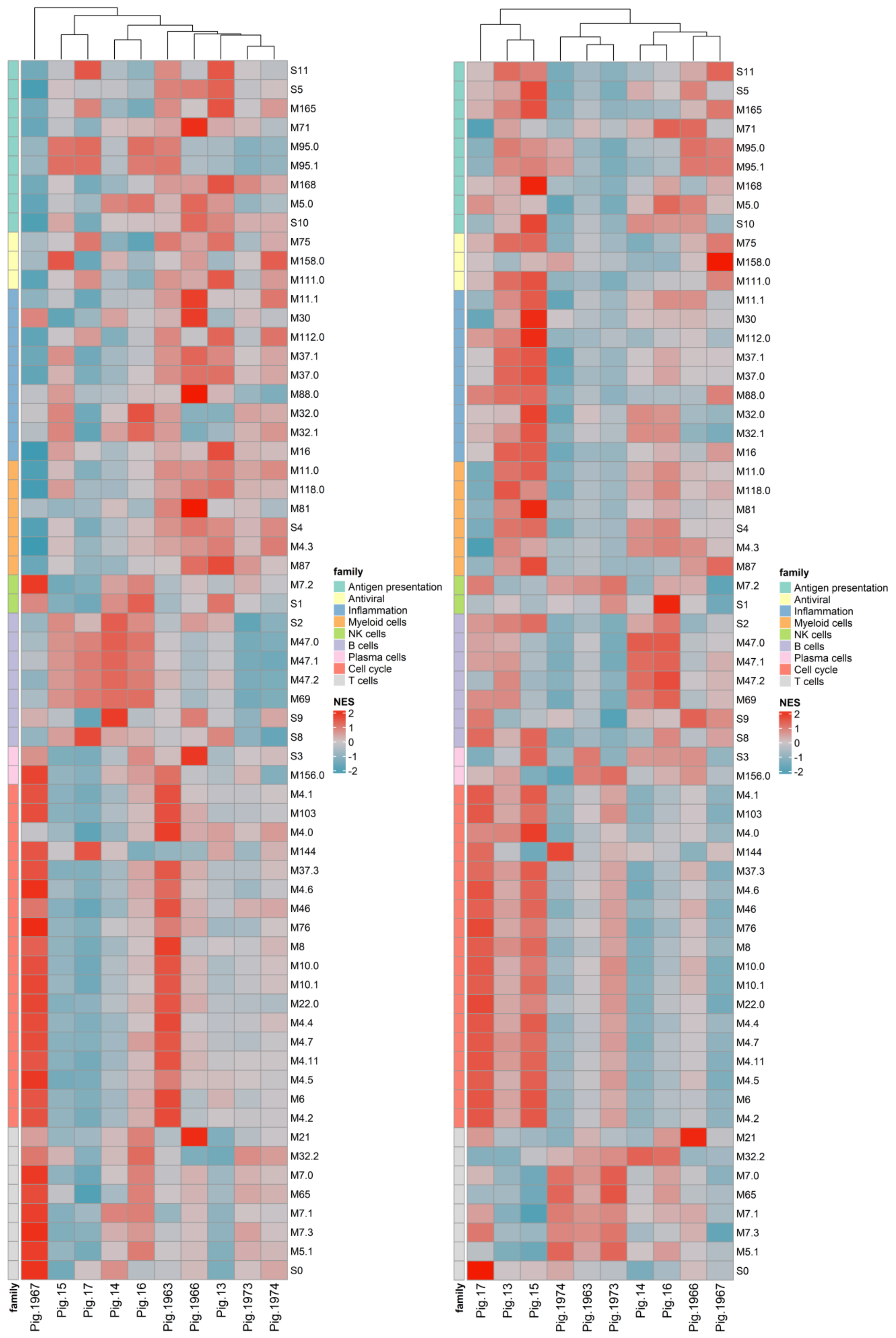


**Supplementary Fig. 21. Heatmaps showing absolute BTM expression in blood leukocytes at 4 dpc (A) and 7 dpc (B).** Gene expression counts were obtained after bulk RNA sequencing and used without normalization to day 0 values. GSVA was then performed with BTM gene sets to calculate NES values displayed on the heatmaps. BTM families are indicated by the color codes (left side) and BTM identifiers are listed on the right side.

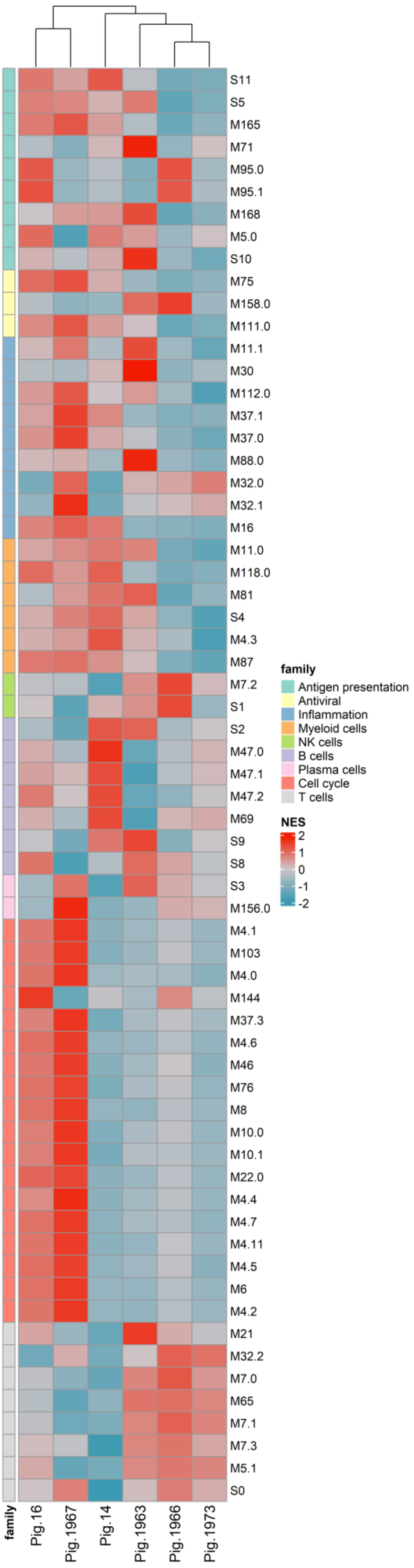


**Supplementary Fig. 22. Heatmap showing absolute BTM expression in blood leukocytes at 11 dpc.** Gene expression counts were obtained after bulk RNA sequencing and used without normalization to day 0 values. GSVA was then performed with BTM gene sets to calculate NES values displayed on the heatmaps. BTM families are indicated by the color codes (left side) and BTM identifiers are listed on the right side.

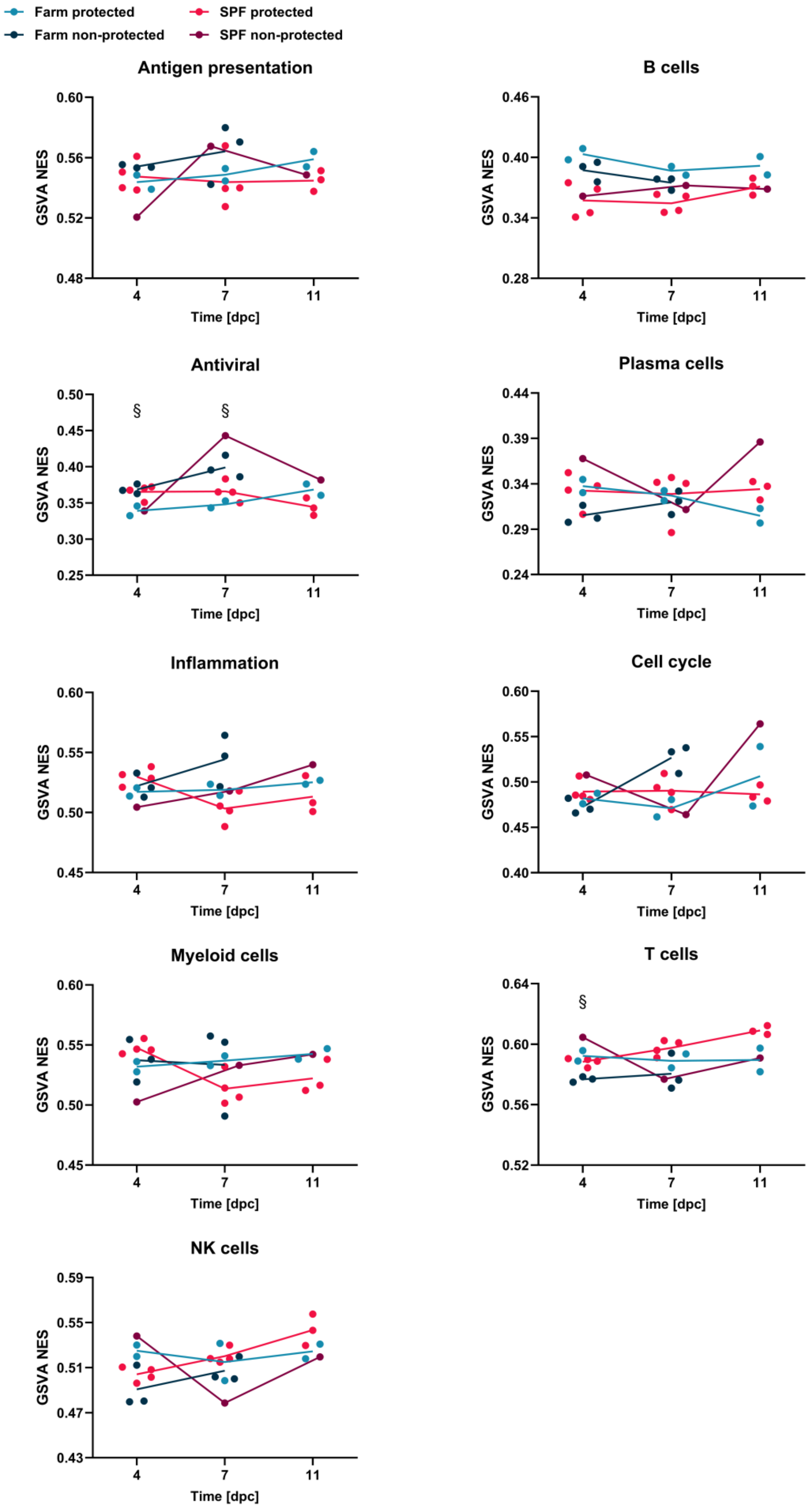


**Supplementary Fig. 23. Kinetics of average NES of BTM families after Armenia 2008 challenge.** NES values for BTM expression from individual animals were calculated by GSVA using non-normalized read counts. At each time point, NES values of BTMs belonging to one family were used to determine average family scores. Plots demonstrate average NES values for each family related to either innate (left column) or adaptive immunity (right column). At 4 and 7 dpc, n = 5 pigs in farm and SPF groups, and at 11 dpc, n = 2 in farm group and n = 4 in SPF group. Differences between protected and non-protected farm pigs were analyzed by unpaired t-test with Holm-Šídák's correction for multiple comparisons; $^{\S}p < 0.05$.
